# Lats1/2 are essential for developmental vascular remodeling and biomechanical adaptation to shear stress

**DOI:** 10.1101/2024.12.01.626284

**Authors:** Mitzy A. Cowdin, Tuli Pramanik, Shelby R. Mohr-Allen, Yuting Fu, Austin Mills, Victor D. Varner, George E. Davis, Ondine Cleaver

## Abstract

Blood vessels in mammalian embryos develop from initial aggregates of endothelial cell (EC) progenitors, which coordinate the opening and stabilization of central vascular lumens, all while under progressively increasing flow and pressure from blood circulation. Mechanical cues exerted by shear stress from the blood flow remodel an initial vascular plexus into a ramifying array of large and small vessels. As plasma starts to fill vascular lumens, these forces trigger changes in EC gene expression and dynamic alterations in cell shape and cell adhesion, as cuboidal angioblasts elongate and flatten into ECs. Little is known about how embryonic ECs sense and transduce hemodynamic signals as vessels form *in vivo*. Here, we report a critical requirement for the Lats1 and Lats2 Hippo pathway kinases during this process. We show that when Lats1/2 are genetically deleted in ECs, embryos develop severe defects in blood vessel formation, which lead to embryonic lethality by E11.5. We find that initial vessel patterning and circulation initiate properly, however remodeling of the initial vascular plexus fails due to lumen collapse and altered blood flow. When Lats1/2 are knocked down using siRNA approaches in cultured ECs, cells fail to elongate and polarize, similar to ECs in the mutant embryos. In addition, VE-cadherin (VEcad) based junctions fail to mature under shear stress. These data show that Lats1/2 deficient cells no longer respond to laminar shear stress, both *in vivo* and *in vitro*. This work identifies the Hippo pathway kinases Lats1 and Lats2 as critical transducers of biomechanical cues during the early steps of blood vessel remodeling. This study will provide new targets for treatment of vascular diseases and new directions for efforts to generate vascularized tissues for replacement therapies.

**Highlights:**

- Lats1 and Lats2 mRNA and protein are expressed in murine embryonic endothelial cells (ECs).
- Deletion of Lats1/2 in embryonic endothelium results in severe vascular defects and embryonic lethality.
- Loss of Lats1/2 leads to failure of both vascular remodeling and EC elongation upon exposure to flow, *in vivo* and *in vitro*.
- Lats1/2 are required for cell-cell VE-cadherin adhesion maturation under flow.
- Loss of Lats1/2 results in cytoskeletal disorganization in response to shear stress.

## Introduction

Blood vessels course through virtually every tissue in our bodies. Building and maintaining them is critical to life. In the early embryo, blood vessels are first built through a process termed *vasculogenesis*, where progenitor cells coalesce and organize to form the first primitive blood vessels.^1,2^ After vasculogenesis, blood vessels grow primarily through a process called *angiogenesis*, where the existing blood vessels undergo rapid expansion via sprouting and remodeling. Remodeling is profoundly influenced by hemodynamic forces. Defects in blood vessel development or integrity can lead to a variety of deleterious consequences including vascular malformations, and stroke.^3,4^

### Biomechanical forces during vascular development

The endothelial cells (ECs) that line our vessels are exquisitely sensitive to physical forces, including shear stress, cyclic stretch, and pressure.^5,6^ Changes in the mechanical environment of ECs are known to drive cell shape changes, stabilization of endothelial junctions, and arteriovenous differentiation.^7–10^ At the forefront of forces that impact ECs is the effect of hemodynamic blood flow. Laminar shear stress (LSS) generated by blood flow drives remodeling and stabilization of vascular lumens after they are formed. Vascular remodeling is the transformation of an initial vascular plexus into a ramifying array of large and small vessels, either by expansion or narrowing of lumens, or by vessel regression.^11^ Vascular lumen expansion is promoted by mechanical signals, such as cyclic stretch and pressure, downstream of cardiac function.^12^ Fluid shear stress caused by blood flow has been shown to be sufficient to drive vascular remodeling in extraembryonic tissues, including the yolk sac.^13^ of endothelial mechanobiology continues to expand.

### Molecular mechanisms of cells sensing biomechanical cues

Biomechanical forces are interpreted by ECs and transduced via cascades of molecular pathways, many of which have been identified in the last few decades.^14^ ECs sense these forces via a complex molecular machinery which includes cell membrane and cytoplasmic molecules that are responsive to biomechanical cues.^6,15,16^ In the early 2000s, the ‘mechanosensory complex’ that dynamically senses shear stress was discovered at endothelial cell-cell junctions, and includes platelet endothelial cell adhesion molecule-1 (PECAM1, or CD31) and vascular endothelial cadherin (VE-cadherin, VEcad).^17^ VEcad junctions dynamically remodel in response to shear stress, where they organize at cell-cell interfaces and ultimately ‘mature’ into tight, linear structures under laminar flow conditions.^18^ Additionally, a variety of mechanosensors are present at the endothelial membrane surface which modulate responses to shear stress and other mechanical factors. Downstream of flow forces, mechanically responsive pathways important during endothelial development including PI3K-Akt, KLF2/4, and Notch signaling pathways which have been linked to endothelial integrity in response to shear stress.^19–22^

More recently identified signaling factors involved in sensing of mechanical cues are members of the Hippo signaling pathway.^23^ In mammals, when Hippo is “on”, the STE20-like protein kinases 1 and 2 (Mst1/2) activate Lats1 and Lats2 (Lats1/2) via phosphorylation. Active Lats1/2 kinases, in turn, phosphorylate the transcriptional coactivators Yes-associated Protein (YAP) and Transcriptional co-activator with PDZ-binding motif (TAZ), leading to their retention in the cytoplasm. When Hippo is “off”, this cascade does not occur and YAP/TAZ translocate to the nucleus to act as transcriptional coactivators, upstream of genes involved in cell proliferation, survival and more. Intriguingly, Hippo signaling is responsive to biomechanical cues, as robust and striking translocation of YAP/TAZ occurs in response to external cues, thereby controlling their transcriptional activities in response to signals such as matrix stiffness or stretch.^23,24^ How upstream Hippo pathway kinases work cooperatively with mechanotransduction to regulate endothelial stability is not well understood.

### Hippo pathway in developing blood vessels

The role of the Hippo effectors YAP/TAZ in blood vessel development has been studied in various contexts. Endothelial-specific YAP/TAZ knockout in embryonic mice lead to developmental angiogenesis defects^25^ and postnatal deletion of YAP/TAZ in ECs results in retinal angiogenesis defects.^26,27^ Additionally, although transcription regulation via YAP/TAZ is thought to be their primary function, the cytoplasmic retention of YAP/TAZ has been shown to affect retinal angiogenesis by controlling Cdc42 activity.^28^ Furthermore, YAP/ TAZ localization is influenced by blood flow. YAP shuttles into the nucleus at the onset of blood flow, but YAP/TAZ translocate to the cytoplasm after sustained laminar flow in zebrafish and cultured ECs.^29,30^

Less is known about the upstream kinases Lats1/2 in the forming vasculature. Lats1/2 have been linked to sprouting angiogenesis during bone angiogenesis.^31,32^ They have also been associated with hyperbranching and proliferation in the retina.^27,28^ Lats1/2 have also been shown to be required in lymphatic vasculature, as YAP/TAZ hyperactivation (caused by Lats1/2 depletion) impairs embryonic lymphangiogenesis and aggravates pathological lymphangiogenesis by suppressing Prox1 expression.^33^ These findings, along with the possibility that Lats1/2 may have other roles beyond facilitating the translocation of YAP/TAZ, argue for the need to study the role of endothelial Lats1/2. Past work leaves an open question in the field of how endothelium is regulated by Hippo signaling in the maintenance of vessel integrity during embryonic development, and additionally, how Hippo signaling may modulate endothelial adaptation to shear stress.

Here, we show that Lats1/2 are critical to embryonic blood vessel lumen stability, endothelial cell shape, and junctional integrity, thereby promoting endothelial resilience. We genetically ablated Lats1/2 specifically in murine ECs at embryonic stages, and this resulted in collapse of endothelial lumens and failure of vascular remodeling. We show that Lats1/2 depleted embryos initiate lumenogenesis and blood circulation normally but develop vessel occlusions by E9.5 and blood flow is blocked. We model these findings in an *in vitro* model of shear stress, and observe that, similar to embryonic findings, Lats1/2 are critical for proper junctional maturation and cell shape changes under shear stress. Additionally, we show the requirement of Lats1/2 for cytoskeletal rearrangements and front-rear polarity under shear stress. Moreover, we demonstrate that Lats1/2 temporally control YAP/TAZ activity under shear stress. Finally, we establish the requirement of YAP/TAZ activity in Lats depletion phenotypes, by genetically co-ablating YAP/TAZ and Lats1/2 in embryos and cultured ECs. Together, this work brings Lats1/2 into the forefront of EC mechanobiology, solidifying the role for canonical Hippo signaling in the maintenance of endothelial stability in embryogenesis and shedding light on a previously under-appreciated role of Lats1/2 in regulating endothelial YAP/TAZ activity under shear stress.

## Methods

### Mouse models

All animal husbandry was performed in accordance with protocols approved by the University of Texas Southwestern Medical Center Institutional Animal Care and Use Committee. Timed pregnancies were set up and noon on the day of visible plug was considered embryonic day (E)0.5. CD1 mice were purchased from Charles River Laboratories and bred for wild type experiments **(Major resources Table S1)**. The following mouse lines were used for relevant experiments: Lats1^flox34^, Lats2^flox 34^, Cdh5-(PAC)CreER^T2,35^ Rosa-CAG-LSL-tdTomato,^36^ TRE-TAZ4SA,^37^ YAP^flox,38^ TAZ^flox,38^ Vecad-tTA^39^ **(Major resources Table S2)**.

For embryonic dissections using the inducible Cdh5-CreER^T2^, tamoxifen was diluted in corn oil and 75 mg/kg body weight was administered to pregnant dams via oral gavage starting at E6.5. For E8.5 and E9.0 dissections, tamoxifen was administered once more at E7.5. For E9.5 and beyond, tamoxifen was administered at E7.5 and E8.5.

### Embryo handling and tissue collection

Dams were euthanized via approved protocols at desired time points and embryos and postnatal tissues were collected and dissected in PBS. Sex was not determined at developmental time points. As developmental time can vary between littermates collected at the same gestational time point, staging was determined by counting somite pairs and/ or visual inspection of developmental landmarks in E8.5-E9.5 embryos. E8.5-E8.75 was classified as 6-14 sp, before (E8.5) or during (E8.75) turning, and before neural fold closure. E9.0 was classified as 15-18 sp, after embryo turning and neural fold closure has occurred. E9.5 was classified as 20-29 sp. Stage-matched embryos were used as controls. Whole embryos were fixed for 30 minutes at room temperature (RT) (E8.5-E9.5) or at 4°C overnight (O/N) (>E9.5) with 4% paraformaldehyde (PFA) in PBS.

### Embryo and yolk sac whole-mount immunofluorescence

Whole-mount staining was performed as previously described with slight modifications (Meadows et al., 2013). Fixed embryos were permeabilized in 1% Triton-X in PBS (PBST) for 1.5 hr then blocked in CAS block (Invitrogen, Carlsbad, CA) for 1.5-2 hr. Embryos were then incubated with indicated antibodies dissolved in CAS block overnight at 4°C. Thereafter, embryos were washed thrice in PBS then incubated with appropriate secondary antibody (1:250, Invitrogen) dissolved in CAS block overnight at 4°C. Antibody information can be found in **Major Resources Tables S3 and S4**. The next day, the embryos were washed in PBS, dehydrated into 100% methanol, then were visualized after clearing in 2:1 benzyl alcohol: benzyl benzoate (BABB). Embryos were mounted in concavity slides in BABB. Images were obtained using Zeiss LSM700 Axio Imager confocal microscope.

### Section IF

For paraffin sectioning, tissues were embedded as described previously with some modifications ^40^. Briefly, tissues were dehydrated to 100% EtOH, washed in xylene, and then incubated in 1:1 xylene: paraffin for 10 minutes at 65°C. After incubating for at least 4 hours at 65°C with paraffin replacement every hour, tissues were embedded in paraffin and sectioned at 10 μm. Paraffin sections were baked at 60°C for 10 minutes and de-paraffinized in xylene (2 x 5 minutes) and rehydrated through EtOH to PBS. All washes were performed using PBS. After washing, tissue sections were permeabilized using 0.3% Triton-X/PBS for 10 minutes. Antigen retrieval was performed under pressure using acidic R-buffer A (Electron Microscopy Sciences) for nuclear antigens and R-buffer B (Electron Microscopy Sciences) for cytoplasmic antigens. Sections were washed, blocked in CAS block for 1 hour (Invitrogen, Carlsbad, CA), and incubated in primary antibodies (diluted in CAS Block, list of antibodies can be found in **Major Resources Table S3** using hybridization chambers O/N at 4°C. The next day, sections were washed and incubated in Alexa Fluor series antibodies (Invitrogen, Carlsbad, CA; 1:500; **Major Resources Table S4**) for 2 hours at room temperature (RT). Sections were then washed and mounted in Fluoromount G with DAPI (ThermoFisher, 00-4959-52), allowed to dry, then stored at 4°C. Images were obtained using Nikon A1R confocal microscope with Nikon software.

### Isolectin perfusion

Dyes such as Evans Blue (1% in PBS) combined with Isolectin B4-488 (1:50 in PBS) (Invitrogen, Carlsbad, CA) were injected into hearts of E9.0 and E9.5 embryos as previously described ^41^ using the World Precision Instruments µPUMP (WPI, PV850) with pulled capillary glass needles. Images were taken shortly after injections with Zeiss Axio Imager M2 epifluorescence microscope.

### Cell culture

Human pulmonary artery endothelial cells (HPAECs) were purchased from Lonza Bioscience and were cultured in Endothelial Growth Medium-2 (cc-3162, Lonza) at 37°C and 5% CO2. ECs within passages 4-8 were used for experiments. Human umbilical vein endothelial cells (HUVECs) were purchased from Lonza and were grown in Supermedia as described (Koh et al, 2008). HUVECs were used from passages 3-6.

### siRNA transfection

HPAECs were seeded at 10,000 cells/ cm^2^ then transfected with indicated siRNA (Dharmacon, ON-TARGET SMARTpool; **Major Resources Table S5**) using Lipofectamine RNAi Max (Invitrogen) the next day according to manufacturer’s instructions. Cells were used for subsequent experiments 48-72 hours post transfection. The efficiencies of gene knockdown were determined by western blot and qPCR analyses.

### Shear stress experiments

Wild type HPAECs were plated onto µ-Slide I Luer, IbiTreat 0.4 mm (Ibidi, 80176) at 25,000 cells/ cm^2^ then exposed to 10 dynes/ cm^2^ the following day for the indicated time periods using the Red Perfusion Set (Ibidi, 10962) or kept as static control. siRNA Transfected ECs were replated onto µ-Slide I Luer, IbiTreat 0.4 mm slides at 15,000 cells/ cm^2^ for long term experiments and 25,000 cells/ cm^2^ for short term (<24 hours) experiments 48 hours post transfection. The following day, HPAECs were exposed to LSS of 10 dynes/ cm^2^ for short term experiments or re-dosed with siRNA to ensure knockdown efficiency for long term experiments. Re-dosed cells were exposed to shear stress the following morning for indicated time points.

### EC tubulogenesis assay in 3D collagen matrices

HUVECs were used for EC tube formation and stabilization assays after addition of the Lats1/2 inhibitor, TRULI (MedChemExpress). EC tube networks were allowed to form for 72 hr in 3D collagen matrices using defined culture media (Medium 199) (Gibco) containing reduced serum-supplement II (which contains insulin) (Koh et al., 2008), as well as recombinant fibroblast growth factor (FGF)-2, stem cell factor (SCF), interleukin-3 (IL-3), and stromal-derived factor-1 alpha (SDF-1α) (Stratman et al., 2011). FGF-2 was obtained from Gibco, while SCF, IL-3 and SDF-1α were obtained from R&D Systems. Media was replaced at 72 hr and varying concentrations of TRULI were added vs. control. After 48 hr, cultures were fixed with 3% glutaraldehyde in PBS and were then stained with 0.1% toluidine blue. EC tube areas were quantitated using Metamorph software, version 7.8 (Molecular Devices).

### Immunofluorescence on cultured ECs

Cultured HPAECs were fixed with 4% PFA, then rinsed with PBS 3 times. Cells were permeabilized in 0.2% Triton-X 100 in PBS for 15 minutes then blocked in CAS block (Invitrogen) for 30 minutes. Cells were then incubated with primary antibodies diluted in CAS block O/N at 4°C (Major Resources Table S4). Afterward, cells were washed in PBS and incubated with secondary antibodies diluted 1:500 in CAS block (**Major Resources Table S4)** for 1 hour, washed in PBS then mounted with Ibidi Mounting medium with DAPI (Ibidi, 50011). Areas with similar confluency were chosen for visualization and analysis to account for different rates of proliferation between cell lots, passages, and treatments. Images were captured using a Nikon A1R Confocal microscope or Nikon CSU-W1 dual camera confocal microscope with Nikon software.

### Protein extraction and Western blot

Cells were rinsed in ice-cold PBS then lysed in RIPA buffer (Abcam) according to manufacturer’s instructions. Protein concentration was determined using the Pierce BCA protein assay kit and concentration of each sample was equalized to that of the lowest concentration using ice-cold RIPA buffer. Subsequent western blot analysis was done as previously described ^40^. Antibody concentrations can be found in supplemental materials.

### RNA extraction and quantitative PCR analysis

Real-time qPCR analysis was done as previously described.^40^ Briefly, RNA was extracted using RNeasy Mini Kit (Qiagen) and cDNA was synthesized using Super-Script III (Invitrogen). qPCR was performed with SYBR Green Master Mix (Applied Biosystems) using gene-specific primers as listed in **Major Resources Table S6.**

### Bulk RNA sequencing

RNA was isolated using the RNeasy Mini Kit (Qiagen) from transfected HPAECs after shear stress experiments by pooling 3-4 Ibidi I Luer slides for each condition. RNA was quantified using a Nanodrop and samples with and evaluated for quality with a bioanalyzer by the UTSW Genomics Core. All samples had a RIN score> 8.

Transcript-level abundance was imported and count and offset matrices generated using the tximport R/Bioconductor package. RNA-seq data have been deposited in the Gene Expression Omnibus (GEO) under the series accession number xxx.

### Embryo length and aorta diameter quantifications

After dissection, embryos were inspected under a stereoscope and embryo length was measured using the maximum total length of each embryo using a sagittal view (crown to rump length, CRL), as previously described^42^. The aortae were detected after immunofluorescence for an endothelial marker (typically PE or Cx40). Dorsal aortae diameters were measured by drawing a straight line across the width of the aorta. Multiple measurements were taken per aorta and the average for each embryo was reported.

### Dorsal midline quantifications

Sagittal whole mount immunofluorescent images of E9.5 embryos stained with PECAM (CD31) and Emcn (PE) were loaded into FIJI (Fiji is just Image J) and maximum intensity projections were created. The edge of the embryo was determined using a separate channel, a 50 μm x 50 μm ROI was created, and intensity of PE was measured within this ROI. 3 ROIs were quantified per embryo along the dorsal midline. Each data point represents an average of 3 individual 50 μm x 50 μm ROI/ embryo normalized to control.

### Cell shape quantifications

Sagittal whole mount immunofluorescent z-stacks of E9.5 embryos stained with VEcad were loaded into FIJI. Optical sections were isolated and VEcad+ cells in the dorsal aortae were manually traced using the freehand selection tool in FIJI. Only cells which had a fully visible “en face” membrane in a sub-stack were measured. At least 15 cells were quantified per embryo. Each data point represents one cell. To quantify nuclear shape in the aortae of embryos, cross sectional paraffin sections stained for PE were loaded into FIJI. Dorsal aortae were identified by anatomy. DAPI+ EC nuclei were identified as cells with PE wrapped around the innermost nucleus touching the lumen of an aorta. The EC nuclei were manually traced using the freehand selection tool in FIJI using DAPI staining and the major and minor axes were measured using Fiji’s measurement tool. At least two sections per embryo were quantified. Each data point represents one nucleus.

### Nuclear YAP percentage quantifications

Measurements of nuclear YAP were determined as previously described ^29^. Briefly, using FIJI, a 7-8 µm circular ROI of consistent size was quantified inside (nuclear) and just outside of the nucleus (cytoplasmic) in one optical section in the middle of the cell based on DAPI staining. The ratio of the Mean Gray value of nuclear YAP intensity was reported as a ratio of nuclear YAP/ cytoplasmic YAP (n/c).

### Golgi polarization quantifications

Maximum intensity projections of ECs were created with FIJI with flow direction going from left to right. Nuclei and Golgi were segmented by creating a binary mask of DAPI and GM130 respectively. Identify Primary Objects was used to segment nuclei and Golgi and positions of each centroid were found using Centroid/ center of mass measurements in Cell Profiler. Angle of each nucleus to nearest Golgi were calculated by measuring angle between the vector of the nucleus centroid point to nearest Golgi centroid point. Golgi polarization was determined by calculating the angle of the nuclear-Golgi vector with respect to the flow direction (images were all rotated if not already oriented so that flow direction was going from left to right to make this calculation consistent). Rose plots of all angles were made using Matlab’s polar histogram function and bins of 15° were used to visualize angles. Angles +/- 45° of flow direction (Golgi in front of nucleus with respect to flow direction, downstream of flow) were classified as “with flow”, +/- 46°-135° are perpendicular, and +/-135°-180° (Golgi behind nucleus with respect to flow direction, upstream) were polarized “against flow”.

### F-actin direction quantifications

To quantify the directionality of F-actin fibers in response to stress, cells were fixed and stained with Phalloidin to visualize F-actin fibers. Corresponding images were loaded into FIJI and directionality distribution was quantified using Directionality plugin in FIJI. Fourier components were used for quantification and 90 bins from −90° to 90° from flow direction created using the plugin. The resultant numbers in each bin were transferred to Prism to create an average histogram; graph is represented as a density plot to better visualize distribution of data along the axis.

### Data analysis and visualization

Data were plotted and analyzed in GraphPad Prism 10. *In vivo* data and quantifications represent at least n=3 embryos from at least 2 separate litters unless stated otherwise in figure legends. *In vitro* data represent at least 3 separate experiments unless otherwise indicated in figure legends. Sample sizes analyzed are provided in figure captions. Both male and female mice were used for experiments. No experiments were randomized for animal studies, and investigators were aware of group allocation for outcome assessment. Statistics were calculated using Prism 10 and statistical tests used are in legends of individual quantifications.\**P*<0.05; \*\**P*<0.01; \*\*\**P*<0.001; \*\*\*\**P*<0.0001. Image J’s de-speckle feature was used in Fig. 4 K-L due to significant background from antibodies. Linear manipulations of images were carried out using FIJI. Figures and models were made using Microsoft PowerPoint and BioRender.com.

## Results

### Loss of Lats kinases in endothelial cells leads to embryonic lethality

To begin our investigation of the role of the Hippo pathway in early vascular development, we first sought to assess expression levels of Lats1 and Lats2 in embryonic endothelial cells (ECs). We first interrogated a published single cell database of mouse embryos shortly after gastrulation^43^ and found that Lats1 and Lats2 (Lats1/2) transcripts are present ubiquitously, with significant expression in the developing endothelium **(Fig. S1A-A”)**. Additionally, we assessed the presence of phosphorylated Lats1 and Lats2 proteins in the early embryo by immunofluorescence (IF) using a previously characterized phospho-Lats1/2 antibody^40^. These assays revealed presence of pLats1/2 in early embryonic ECs (E8.75), as well as most surrounding embryonic cells (**Fig. S1B**).

To test the role of Lats kinases in early endothelial development, we generated an inducible endothelial-specific *Lats1* and *Lats2* knockout mouse. This was accomplished by mating *Lats1^ff^;Lats2^ff^*females with *Lats1^ff^;Lats2^ff^;Cdh5CreER^T2^*males to yield *Lats1^ff^;Lats2^ff^;Cdh5CreER^T2^*offspring (or *Lats^iECDKO^*) (**Fig. 1A**). To induce deletion of *Lats1* and *Lats2* in endothelial progenitors, or angioblasts, at a time when the first vessels were forming, we administered tamoxifen to timed-pregnant females daily starting at embryonic day (E) 6.5. This approach caused embryonic lethality between E10.5 and E11.5 (**Table 1**).

**Figure 1.**
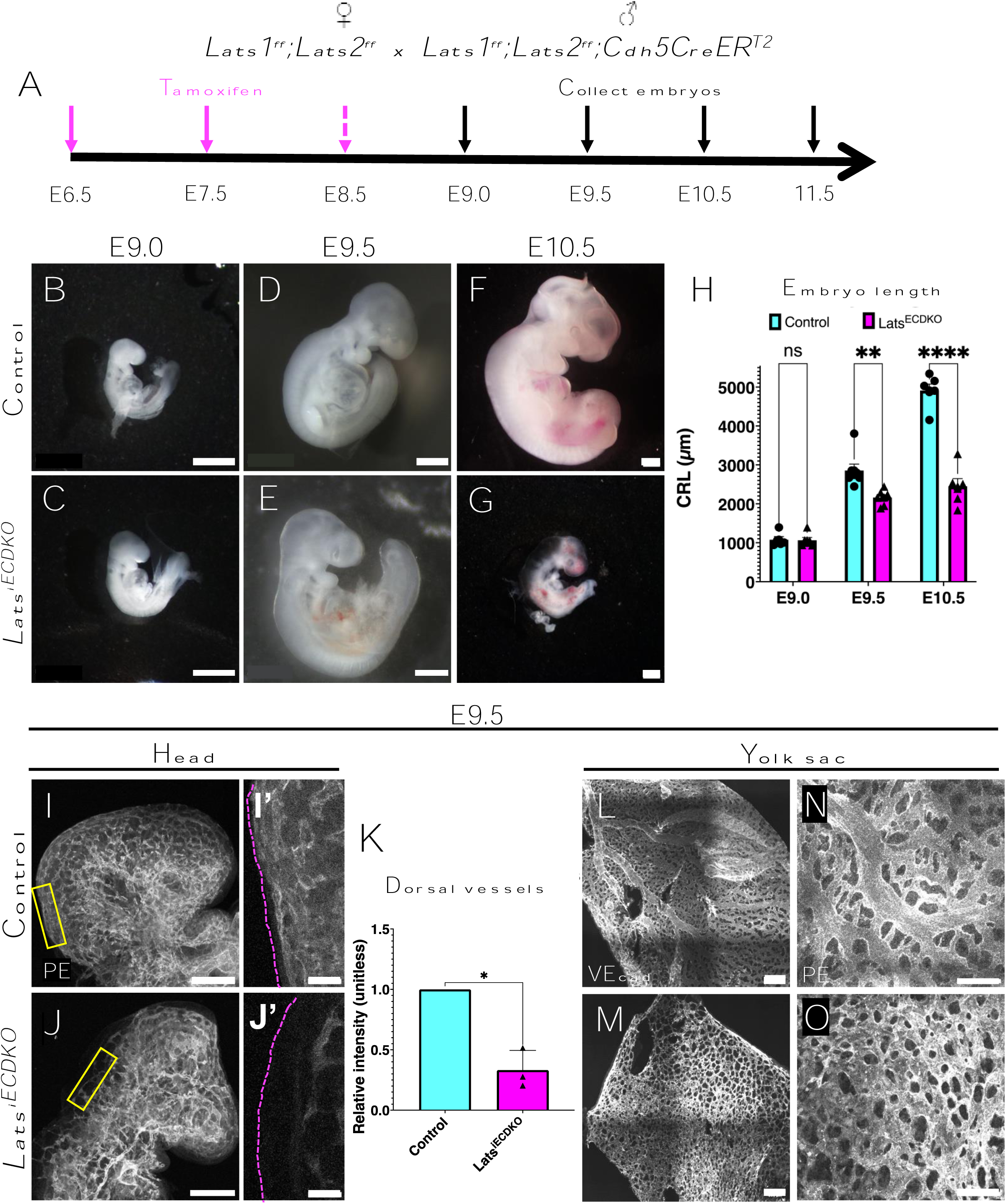
Depletion of EC-specific Lats1 and Lats2 results in embryonic lethality and remodeling defects. **(A)** Tamoxifen scheme for *Lats^iECDKO^* genetic deletion in embryonic endothelium. Pink arrows indicate time of gavage. Stippled arrow represents a time of gavage for collections past E9.0. Black arrows represent timing of collection of embryos. Representative brightfield images of gross phenotype of control (**B, D, F**), and *Lats^iECDKO^* (**C, E, G**) embryos at E9.0 (**B, C**), E9.5 (**D, E**), and E10.5 (**F, G**). Scale bars= 500 µm. **(H)** Quantification of embryo crown to rump length (CRL) of control and *Lats^iECDKO^* embryos at E9.0, E9.5, and E10.5. n=5-6 embryos/ genotype for E9.0, n=7 embryos/ genotype for E9.5, and n=6 embryos/ genotype for E10.5. **(I, J)** Representative whole mount immunofluorescent images of control (I) and *Lats^iECDKO^* (J) vessels in head region. **(I’, J’)** Enlarged insets of boxed regions in I, J showing dorsal vessels in control (**I’**) and *Lats^iECDKO^* (**J’**). **(K)** Quantification of intensity in dorsal vessels of *Lats^iECDKO^* heads normalized to control vessels (n= 3 embryos/ genotype, unitless). **(L-O)** Representative stitched images of VEcad (**L, M**) and CD31/ Endomucin (PE) (**N, O**) immunostaining in control and *Lats^iECDKO^* yolk sacs at E9.5 (scale bar= 200 µm (L,M), 50 µm (N, O). n > 3 embryos/ genotype). Data are shown as mean =/- SEM. Statistical significance determined by Two-way ANOVA with multiple comparisons (H) or unpaired t-test (two-tailed) (K). ns= not significant, * indicates p-value<0.05, ** indicates p-value <0.01, **** indicates p-value< 0.0001.

**Table 1.**
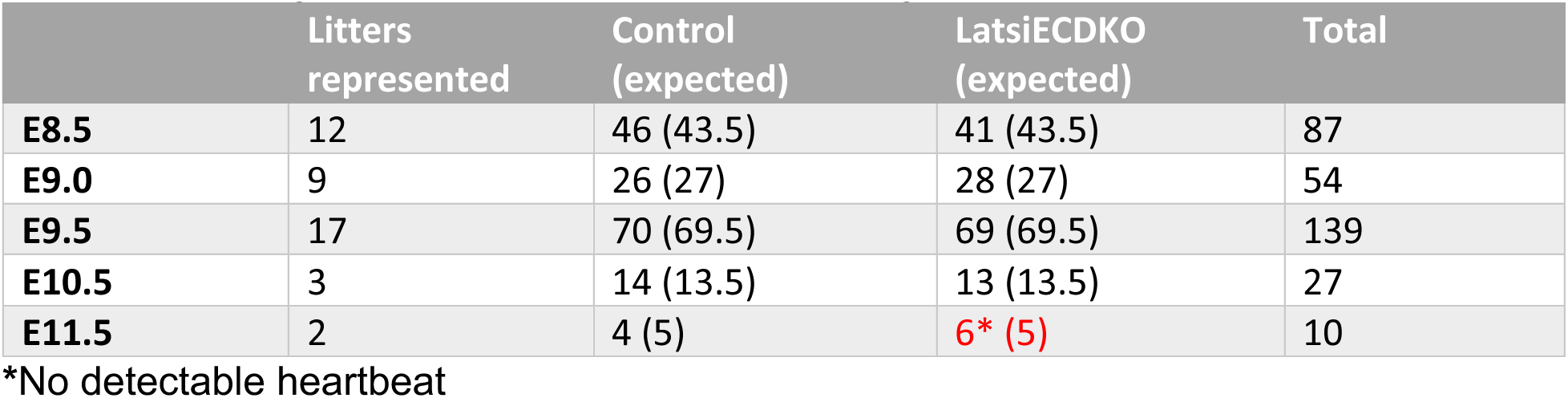
Viability of control and *Lats^iECDKO^* embryos.

|  | Litters represented | Control (expected) | LatsiECDKO (expected) | Total |
| --- | --- | --- | --- | --- |
| **E8.5** | 12 | 46 (43.5) | 41 (43.5) | 87 |
| **E9.0** | 9 | 26 (27) | 28 (27) | 54 |
| **E9.5** | 17 | 70 (69.5) | 69 (69.5) | 139 |
| **E10.5** | 3 | 14 (13.5) | 13 (13.5) | 27 |
| **E11.5** | 2 | 4 (5) | 6* (5) | 10 |

\*No detectable heartbeat

To analyze possible defects in emerging vessels, we characterized control and mutant embryos, from E8.5 to E11.5 (**Fig. 1B-G, Fig. S2A-D**). At E11.5 Cre-negative *Lats1^ff^;Lats2^ff^* (control) embryos displayed normal development (**Fig. S2A**) and a strong fetal heartbeat (**Supplementary video 1**). By contrast, *Lats1^ff^;Lats2^ff^;Cdh5CreERT2* (*Lats^iECDKO^*) embryos at this stage exhibited no discernable heartbeat, were smaller and appeared white and opaque (**Fig. S2B and Supplementary video 2**). To address the cause of lethality in *Lats^iECDKO^*embryos, we analyzed earlier stages. We hypothesized lethality could be due to a defect in *de novo vasculogenesis,* which initiates between E8.0 and E8.25^2^. However, at E8.5, *Lats^iECDKO^* embryos appeared grossly normal when compared to control littermates (**Fig. S2B**). Similarly, at E9.0, mutant and control embryos were similar in terms of overall embryo size **(Fig. 1B, C)**, measured by crown to rump length **(Fig. 1H)**. By E9.5, however, *Lats^iECDKO^* embryos were slightly but significantly smaller than stage-matched controls, despite somite counts being equal (Fig. 1D, E, Fig. S2E). Additionally, by E10.5, *Lats^iECDKO^* embryos displayed severe hemorrhage, pericardial effusion, and edema within the embryo proper, indicative of poor cardiovascular function (**Fig. 1G**) and were significantly smaller than control embryos **(Fig. 1F, H)**. Evaluating *Lats^iECDKO^*;*tdTomato^ff^*embryos, we found tdTomato expression was present in *Lats^iECDKO^*by E8.5, (**Fig. S2F**). This indicates that recombination has occurred by this stage and that vascular defects are observed between 24 and 48 hours thereafter.

We next assessed arterial specification in the early vasculature, at a timepoint when recombination has occurred. We previously showed that arterial gene expression initiates shortly after lumenogenesis^1^, as hemodynamic flow induces remodeling of the initial vascular plexus. We tested for expression of arterial Neuropilin-1 (Nrp1) in the early vasculature and found its expression indistinguishable in E8.5 mutant and control embryos (**Fig. S2G-H’**). Together these data indicate that Lats1/2 are required for vascular development and embryonic survival, but the initial steps of vasculogenesis, including patterning of the initial plexus and arterial specification, occur normally using this model system.

### Endothelial Lats1/2 deletion results in vascular remodeling defects

Given the growth arrest evident between E9.5 and E10.5, a time when vascular remodeling is known to be actively occurring,^13^ we characterized the architecture of *Lats^iECDKO^* developing blood vessels. During this time, the initial net-like vascular plexus transforms into a hierarchical and ramifying array of large and small vessels. To assess this process in *Lats^iECDKO^* embryos, we performed whole mount IF staining for the endothelial markers PECAM1 (CD31) and Endomucin (double stained, hereby designated as “PE”) at E9.5. We found severe defects in both large and small vessels throughout embryonic tissues in *Lats^iECDKO^*compared to control littermates (**Fig. 1I, J, Fig. S2I-J**). We observed differences in the density and morphology of vessels in the head **(Fig. 1I, J)** and trunk region of the embryo **(Fig. S2I, J).** Control embryos had a well-formed vascular tree in the head, comprising of a hierarchical vascular network that extended to dorsal midline of the embryo **(Fig. 1I’)**. By contrast, while a plexus structure was present in *Lats^iECDKO^* embryos at this stage, there was little to no hierarchy in the vascular structures and the branches did not reach the dorsal most midline **(Fig. 1J’, K)**.

We then assessed the yolk sac vasculature which develops concomitant with the formation of the embryo proper.^11,44,45^ IF for PE revealed that yolk sac vessels from control embryos at E9.5 displayed the classical remodeled hierarchical vessel structure, with large vessels ramifying into smaller capillary branches (**Fig. 1L, N**). This transformation has been associated with establishment of blood circulation and shear stress, which leads to remodeling of the vascular architecture.^13^ By contrast, we found that *Lats^iECDKO^* yolk sac vessels at E9.5 remained a primitive plexus, with a honeycomb-like structure and an absence of a hierarchical structure (**Fig. 1M, O**). These data suggest a critical requirement for Lats1/2 during remodeling angiogenesis.

### Lats1/2 are required for DA lumen diameter and stability

The failure of the vascular plexus to remodel into a vascular tree in *Lats^iECDKO^* embryos could be due to failure of aortic lumen formation.^2,46,47^ Therefore, we assessed the lumen of the aortae in control and *Lats^iECDKO^* embryos at various embryonic stages **(Fig. 2A-E)**. Although the aortic width was initially similar between control (mean= 44.86 µm) and *Lats^iECDKO^* (mean= 46.53 µm) embryos at E9.0 **(Fig. 2A-B, E)**, by E9.5 the *Lats^iECDKO^* embryos displayed markedly more narrow lumens (mean= 34.53 µm compared to 79.33 µm), at all anteroposterior locations along the length of the embryonic axis (**Fig. 2D, E**).

**Figure 2.**
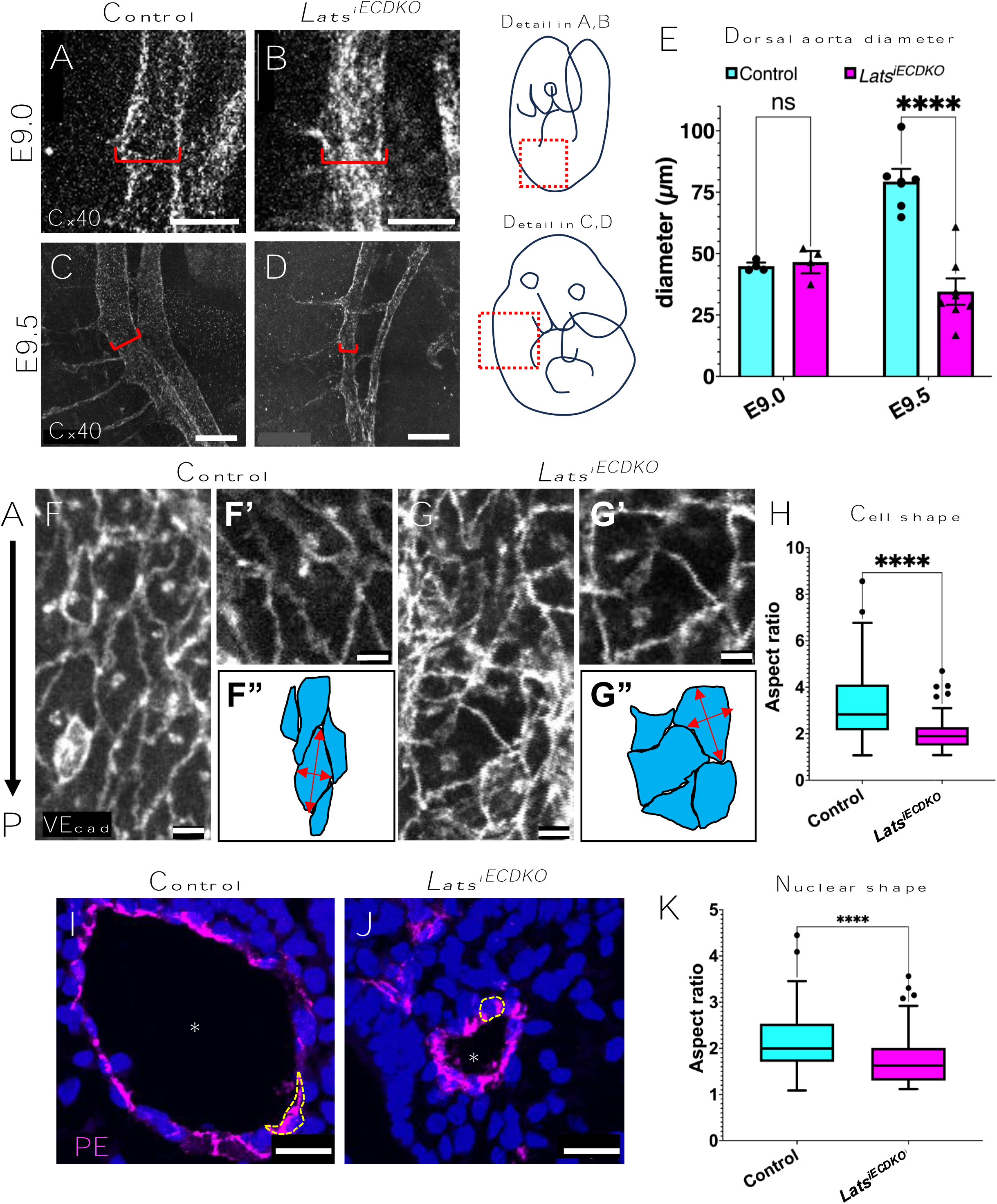
Depletion of endothelial Lats1/2 results in aortic constrictions and cell shape defects. **(A-D)** Vessels labelled with Connexin 40 (Cx40) protein in E9.0 (**A, B**) and E9.5 (**C, D**) aortas in control (**A, C**) and *Lats^iECDKO^* (**B, D**) embryos. Red brackets indicate diameter of aortae. Embryo schematics on right indicate detail view in A, B (top) and C, D (bottom) respectively. Scale bar= 50 µm (A, B), 500 µm (C, D). **(E)** Quantification bar graph of aorta diameter at E9.0 (n=3 embryos/ genotype) and E9.5 (n= 6-7 embryos/ genotype). Each dot represents an individual embryo; error bars represent SEM. **(F-G)** Optical sub-stack of immunostaining for VE-Cadherin (VEcad) in control and mutant embryonic aortae. Plane of optical sections along aorta is sagittal and images represent *en face* view of endothelium (A= anterior; P= posterior). Scale bar= 5 µm. (**F’, G’**) Enlarged view of aortic ECs in control (**F**) and mutant (**G**) embryo aortic ECs. (**F’’, G’’**) Diagram of EC outlines showing differences in cell shapes between control (**F’**’) and mutant (**G’**’) embryo aortic ECs. Red arrows indicate estimated major and minor axes used for quantifications. **(H)** Tukey box and whisker plot of aspect ratio (major axis/ minor axis) of control and *Lats^iECDKO^* aortic ECs. Each value represents an individual cell; n= 3 embryos/ genotype. **(I, J)** Confocal images of transverse paraffin sections through dorsal aortae stained for PE (magenta) and DAPI (blue). Asterisk represents dorsal aorta lumen. Dashed line indicates outline of EC nucleus. Scale bar= 20 µm (**K**) Tukey box and whisker plot of aspect ratio of control and *Lats^iECDKO^* aortic ECs. Each value represents an individual nucleus; n=3 embryos/ genotype. Statistical significance determined by Mann-Whitney U test (H, K). *** indicates p-value <0.001, **** indicates p-value< 0.0001.

### Lats1/2 loss leads to modest reduction of EC proliferation but not apoptosis

In assessing vessels in the *Lats^iECDKO^* mutants, we had noted that the dorsal aortae were greatly reduced in diameter compared to controls at E9.5 **(Fig. 2B, D, E)**. Cross sections of aortae in E9.5 embryos displayed slightly fewer EC nuclei compared to control embryos (control mean= ∼10 nuclei, *Lats^iECDKO^* mean= ∼8) **(Fig. S3A).** The modest decrease in aortic ECs at E9.5, however, does not fully account for the ∼1/2 reduction of aortic lumen diameter in *Lats^iECDKO^*embryos. We next asked whether failed aorta lumen expansion in *Lats^iECDKO^* embryos was due to loss of ECs via cell death in the absence of Lats1/2. We found that apoptosis, quantified by IF for cleaved caspase 3 (cc3), was low and not significantly different in control and *Lats^iECDKO^* embryo ECs **(Fig. S3B-D),** suggesting that cell death did not cause vessel failure at this stage.

We next asked whether there was a cell-intrinsic difference in proliferative capacity or apoptosis in ECs depleted of Lats1/2. Therefore, to model our *in vivo* findings, we treated human pulmonary artery endothelial cells (HPAECs) with siRNA for Lats1 and Lats2 concurrently (siLats1/2). Treatment with siLats1/2 resulted in a dramatic loss of Lats1 and Lats2 transcripts 48 hours after transfection **(Fig. S4A)**. Additionally, protein levels of Lats1 and Lats2 were reduced 72 hours post transfection **(Fig. S4B)**. Importantly, YAP staining was visibly higher in Lats1/2 depleted cells **(Fig. S4C)**, validating our model. We then performed IF for phospho-histone 3 (pHH3) to measure proliferation and cc3 in the Lats1/2 knockdown and control cells. Surprisingly, we found that ECs *in vivo* in the *Lats^iECDKO^,* reduction of Lats1/2 by siRNA (siLats1/2) did not alter total levels of pHH3 at 48-, 72-, or 96-hours post transfection **(Fig. S5D-E).** Additionally, cultured ECs had low cc3 staining and treatment with siRNA for Lats1/2 did not alter total levels of cc3 staining area at 48-, 72-, or 96-hours post transfection **(Fig. S5F-G).**

To further test the contribution of Lats1/2 to lumen maintenance and stability of endothelial tubes following their formation, we seeded HUVECs in 3D collagen matrices for 72 hours to allow for the formation of endothelial tube networks. We have previously demonstrated that ECs form stable vascular tubes in this model.^48^ After allowing tubes to form for 48 hours, we subsequently treated the networks with the Lats1/2 inhibitor TRULI at varying concentrations.^48,49^ After 48 hours of drug treatment, TRULI treatment led to collapse of lumens in this pre-existing network, whereas untreated ECs maintained their lumens **(Fig. S4H)**. This effect was dose dependent **(Fig. S4I)**. This led us to hypothesize that Lats1/2 activity may be required for maintenance of vascular lumens.

### Lats1/2 are required for aortic cell shape

To gain a better understanding as to why the lumens were not maintained in our mutant embryos, we turned our attention to the cell shape, as it is a cellular feature that dramatically changes between lumenogenesis and lumen maintenance.^8,50^ We observed ECs in aortae of *Lats^iECDKO^*embryos exhibited a markedly different cell shape compared to control embryos. Using VEcad IF to outline EC boundaries, we observed a reduced elongation of *Lats^iECDKO^* aortic ECs at E9.5 along the anterior-posterior axis (**Fig. 2F-G”**). The dimensions of individual aortic ECs were quantified by their aspect ratio, using the ratio of the major axis to minor axis of the ECs in optical sections and verified that indeed, individual aortic ECs in *Lats^iECDKO^* were less elongated (median= 1.894 compared to control median= 2.833) **(Fig. 2H).** Additionally, transverse cross sections of E9.5 embryos through the dorsal aortae also revealed ECs were strikingly cuboidal within the *Lats^iECDKO^*aortae, compared to the more flattened ECs of control embryo aortae **(Fig. 2I-J)**. As the cells change shape after lumenogenesis, endothelial cell nuclei become more flattened^50^. Therefore, we measured the ratio of the major and minor axis of the EC nuclei in transverse sections of the dorsal aorta and found that the *Lats^iECDKO^*aortic nuclei do not flatten to the same extent as nuclei in the control embryos **(Fig. 2K).** These data show that Lats1/2 are required for control of EC shape changes during blood vessel morphogenesis, likely allowing the proper expansion of the dorsal aortae diameter.

### VE-cadherin junctions are dysregulated in *Lats^iECDKO^* embryos

Redistribution of junctions to organize into stable cell-cell contacts is critically important for stability of ECs during development and response to external stimuli, including the shear stress imposed by blood flow.^9,51,52^ Therefore, we hypothesized that Lats1/2 may regulate endothelial junctions. We assessed VEcad junctions between ECs in control and *Lats^iECDKO^*embryos at E9.5 and found that VEcad both displayed discontinuous regions as well thicker junctions between the ECs of *Lats^iECDKO^* aortae (**Fig. 3A-B’**). These junction defects were also evident in the yolk sac vessels at E9.5 **(Fig. 3C, D**). To determine when this failure begins to occur, we assayed yolk sac vessels at E9.0, a time where blood flow is known to begin remodeling these vessels. At E9.0, we found initiation of accumulations of VEcad in capillary plexus ECs in *Lats^iECDKO^* yolk sacs **(Fig. 3E-F’**) compared to continuous junctions already present in control vessels. These data suggest that ECs in *Lats^iECDKO^* embryos struggle to mature VEcad based junctions and, as a result, may not be able to fully respond to the cues that drive EC elongation and vessel remodeling.

**Figure 3.**
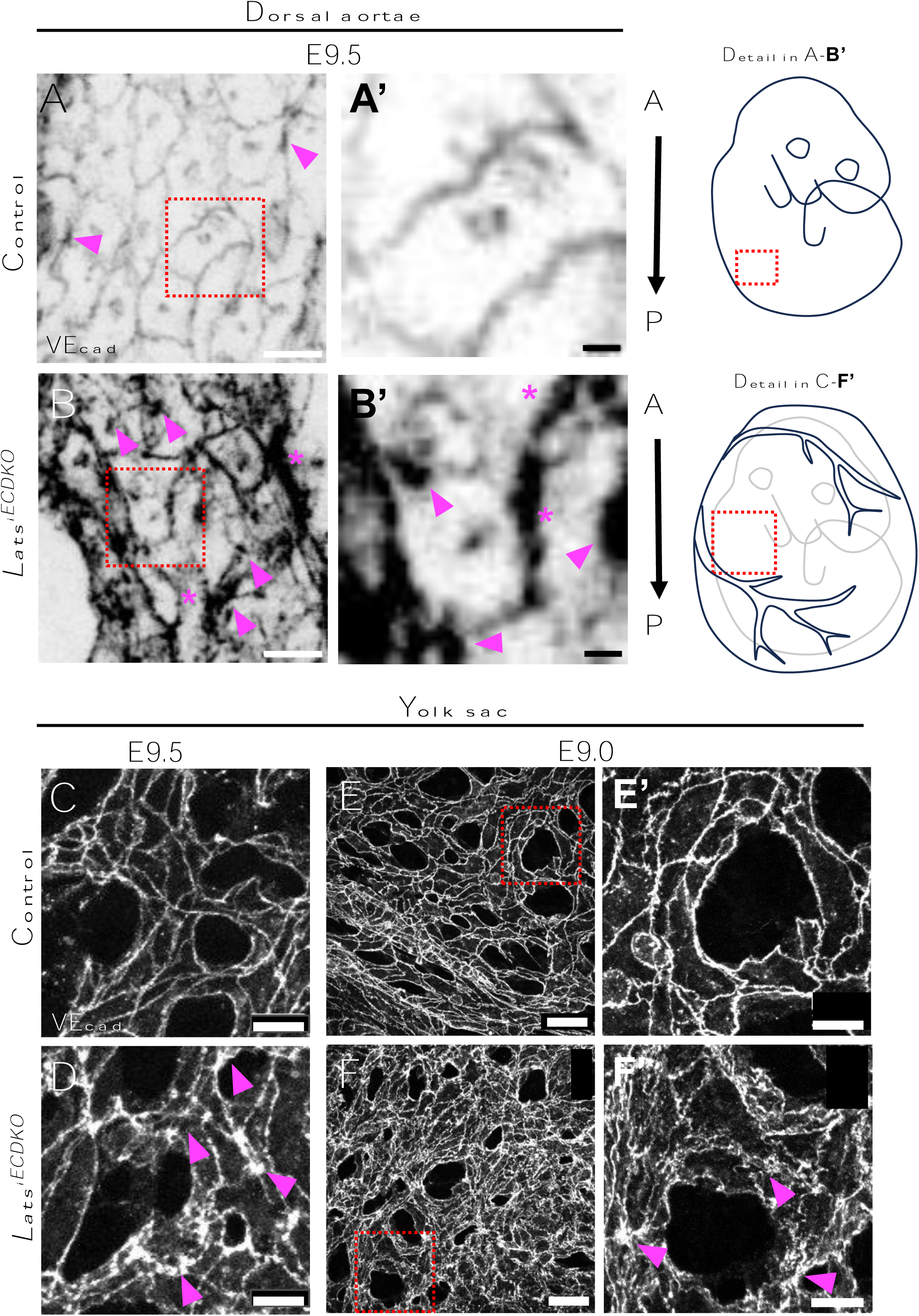
Loss of endothelial Lats1/2 results in altered adherens junction morphology in the developing vasculature. **(A-B)** E9.5 dorsal aortae labeled with VEcad showing altered junction morphology in *Lats^iECDKO^* embryos. Arrowheads indicate areas VEcad aggregates. Asterisks indicate regions of thickened junctions. Scale bar= 10 µm. (**A’,B’**) Magnification of boxed regions in A and B respectively. Scale bar= 2 µm. A= anterior; P= posterior. Embryo schematics on right indicate detail view in A, B (top) and C, D (bottom) respectively. **(C-F)** Representative confocal z-projections of whole mount yolk sac stained for VEcad at E9.5 (**C, D**) and E9.0 **(E-F’).** Arrowheads indicate areas where VEcad has accumulated. Scale bar= 50 µm. **(E’, F’)** magnification of boxed regions in E and F respectively. Scale bar= 20 µm.

### Circulation initiates normally in *Lats^iECDKO^* embryos but fails by E9.5

The defects seen in the *Lats^iECDKO^* embryos suggested cardiac failure and/or compromised circulation. To determine when and if blood flow was impaired, and if it occurred before the phenotypes we observed, we injected isolectinB4-488 (IB4) combined with Evans Blue into the heart of E9.0 and E9.5 embryos. At E9.0, we observed the injected IB4 rapidly traveling through the pumping heart into the aortae in control and mutant embryos **(Fig. 4A-B’, C, D)**. However, starting at E9.5, while the control embryo hearts circulated the fluorescent dye within the branchial arch arteries and dorsal aortae properly **(Fig. 4E, E’, G)**, the *Lats^iECDKO^* embryos failed to pump the dye from the heart into these major vessels **(Fig. 4F, F’, H).** We hypothesized that the failed circulation was due to occlusions or emergence of discontinuities in the vessels in the embryos, leading to loss of blood flow.

**Figure 4.**
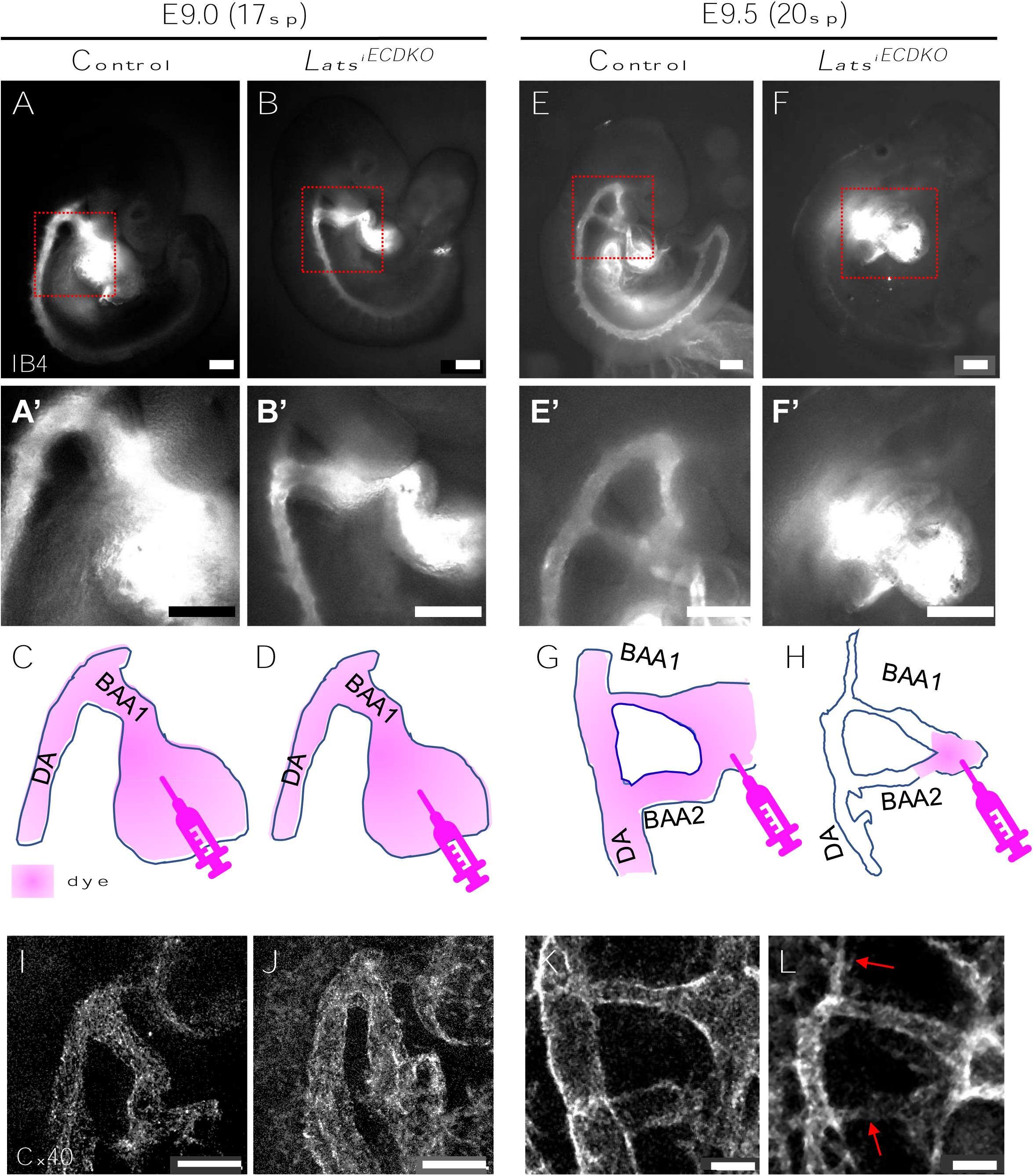
*Lats^iECDKO^* embryos initially establish circulation that later fails due to lumen occlusions. **(A-D)** Circulated Isolectin B4 (IB4) in E9.0 and E9.5 embryos after injection into heart. IB4 passage was observed into the dorsal aortae of both control and *Lats^iECDKO^* embryos at E9.0, making its way down to the embryonic tail (**A, B, C, D**). At E9.5, ink was observed in both the branchial arch arteries (BAA) and dorsal aorta of control embryos (**E, E’, G**), however it was notably absent in *Lats^iECDKO^* mutant embryos at E9.5 (**F, F’, H**). (**A’, B’, E’, F’**) Higher magnifications of boxed regions in A, B, E, F respectively. **(C, D, G, H)** Cartoons represent diameter and orientation of vessels in each condition. Pink represents IB4 passage after injection. **(I-L)** Whole mount immunofluorescence showing normal, uniform caliber vessels in all embryos except E9.5 *Lats^iECDKO^* embryo (**L**), which has narrow lumens in BAA and in aorta. Arrows pointing to narrow lumens. Scale bar= 100 µm. sp= somite pairs.

Analysis of the branchial arch arteries (BAA), which connect the dorsal aorta to the heart, using staining for showed that this critical vessel was established normally in *Lats^iECDKO^* embryos at E9.0 compared to controls, at a time when flow is occurring **(Fig. 4I, J)**. However, by E9.5, the lumens of the BAA and dorsal aorta were collapsed. Control BAA and aortas were open and continuous **(Fig. 4K)**, however, *Lats^iECDKO^*embryos had narrower BAAs leading to an aorta with a significantly reduced luminal space **(Fig. 4L**, arrows). All together, these data suggest that vasculogenesis of the dorsal aortae and large vessels in mutants occurs normally, lumens open, and blood flow is established, but the stability of the blood vessels in *Lats^iECDKO^*embryos is compromised, leading to rapid collapse of vessels, ultimately leading to failure of vascular remodeling and to arrested development.

### Endothelial elongation and Golgi polarization under flow require Lats1/2

Our studies thus far have shown that *Lats^iECDKO^* embryos initially establish a normal vasculature and robust blood flow, but that remodeling angiogenesis and vessel integrity fail shortly thereafter. We note with interest that endothelial failures initiate as circulation ramps up. Given that junctions progressively display defects after onset of blood flow, even while blood flow is still present, we hypothesized that Lats1/2 depleted ECs might fail to respond to mechanical cues from blood flow that are necessary for normal blood vessel morphogenesis. The effect of shear stress is the most prominent and well-studied mechanical cue on embryonic vascular morphogenesis.^11,13,16,53–55^ To test whether Lats1/2 are indeed required for endothelial response to shear stress, we subjected HPAECs to a physiologically relevant laminar shear stress (LSS) of 10 dynes/ cm^2^.^56^ Shear stress exposure for 48 hours, as expected, caused control, non-targeting siRNA-treated ECs (siNT) to elongate (**Fig. 5A-B’**), However, the siLats1/2 ECs under flow had an attenuated elongation (**Fig. 5C-D’**). This was quantified by measuring the aspect ratio of control and siLats1/2 cells with and without shear stress **(Fig. 5E),** which revealed a much lower aspect ratio of siLats1/2 cells under flow compared to control ECs (mean= 1.994 +/- 1.004 compared to siNT flow mean= 5.004 +/- 2.377, p-value< 0.0001).

**Figure 5.**
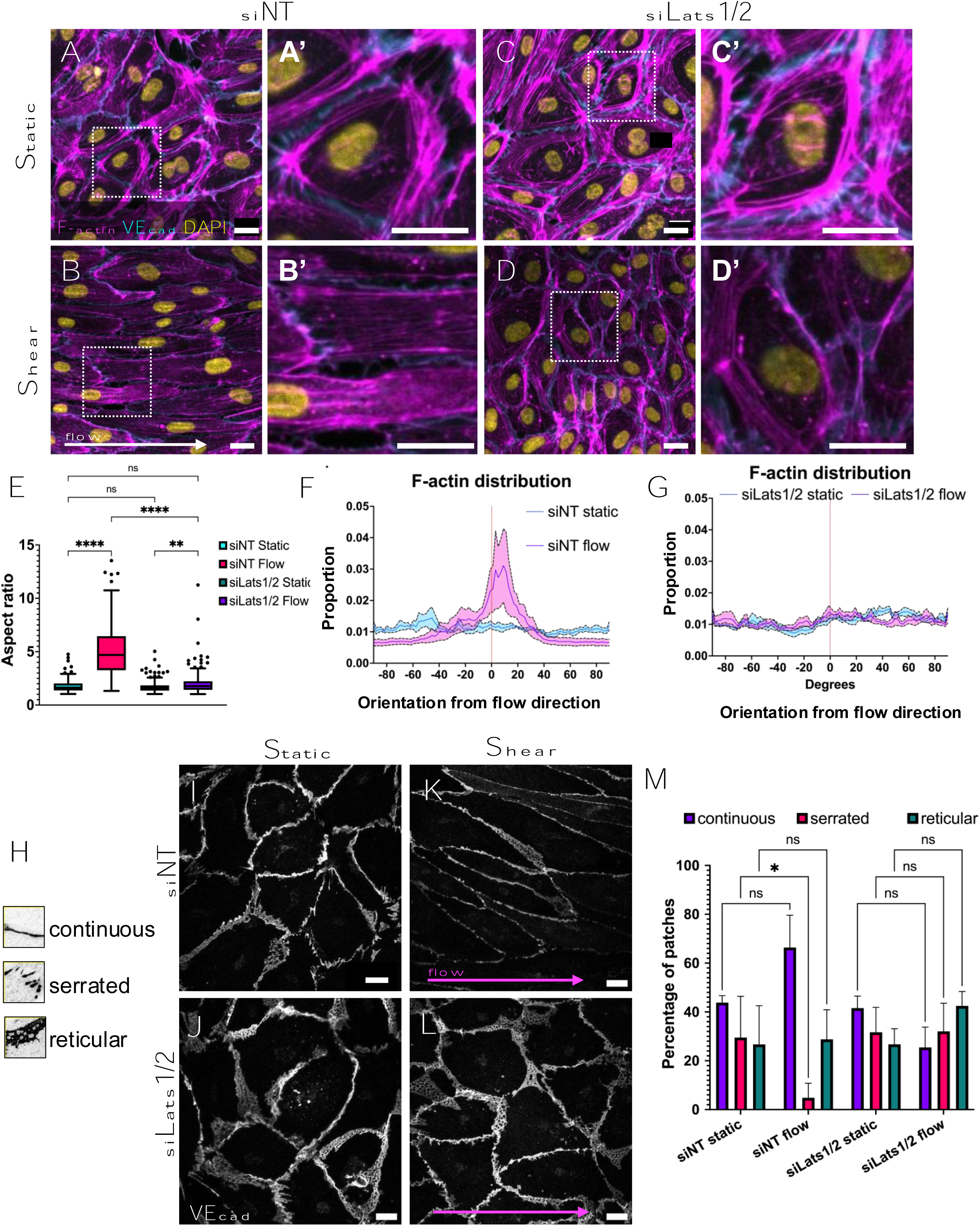
Lats1/2 regulate cell shape, cytoskeletal rearrangements, and junctional maturation under shear stress. **(A-D)** Representative images of stress fiber formation (Phalloidin, magenta; VEcad, cyan; DAPI, yellow) in cells treated with siNT and siLats1/2 in static (**A, A’, C, C’**) and shear stress (**B, B’, D, D’**) conditions. White arrow represents flow direction. **(A’, B’, C’, D’)** Higher magnification views of red boxed areas in A-D respectively **(E)** Quantification of elongation of cells from experiments in A-D. Each value represents an individual cell; n= 3 replicates; at least 200 cells were analyzed per condition. **(F, G)** Quantifications of distribution of F-actin stress fiber alignment with respect to flow direction in siNT (**F**) and siLats1/2 (**G**). Note that siNT cells under flow (pink line, E), have a strong peak within 30 degrees from axis, while siLats1/2 cells do not exhibit a strong peak along the line. Solid line represents average density plot and shaded region represents S.E.M. n= 3 replicates. **(H)** Representative details used for manual classification in 3 categories: continuous, serrated, and reticular. Images analyzed are (I-L) and quantification shown in (M). **(I-L)** Immunofluorescence images of VEcad in control cells (**A, D**) and cells knocked down for Lats1/2 (**B, D**) under static (**I, J**) and shear stress (**K, L**) conditions. Pink arrow represents direction of flow. Scale bar= 20 µm. **(M)** Morphological analysis of VEcad in control and siLats1/2 cells in static and flow conditions. Bars represent percentage of patches analyzed in each category; sum of patches equals 100% in each experimental condition. Error bars represent SD. n= 3 replicates; at least 90 VEcad junctions were analyzed per condition. Statistical significance determined by Kruskill-Wallis test with multiple comparisons (E) or Two-way ANOVA with multiple comparisons (K). ns= not significant, * indicates p-value<0.05.

The alignment of ECs in response to shear stress is thought to be largely mediated by the actin cytoskeleton.^18^ Therefore, we examined whether the actin cytoskeleton responded normally to shear stress the absence of Lats1/2. We examined F-actin orientation using Phalloidin, which revealed that although both control and siLats1/2 cells had cortical actin bundles in static conditions **(Fig. 5A’, C’)**. Shear stress induced reorientation and reorganization of F-actin stress fibers to span the length of ECs parallel to the direction of shear stress in control cells (**Fig. 5B’**); however, Lats1/2 knockdown cells showed no actin reorganization; instead, the cells retained cortical bundles which did not align in any particular direction (**Fig. 5D’**). We quantified cytoskeletal alterations by measuring the direction in which stress fibers aligned along the direction of shear stress using the Directionality tool in FIJI. Fibers in both control and siLats1/2 cells displayed a relatively random alignment under static conditions. Under shear stress, control cells exhibited a peak of stress fibers oriented along the direction of flow (within +/- 45°) **(Fig. 5F)**; however, F-actin directionality in siLats1/2 cells remained relatively unaffected under shear stress conditions **(Fig. 5G)**.

Another measure of ECs sensing hemodynamic forces is Golgi polarization along the axis of the vessel. Golgi localization shifts upstream of the nucleus with respect to shear stress, both *in vivo* and *in vitro*.^57^ We predicted that if Lats1/2 played a role in adapting to flow, siLats1/2 cells would not properly orient their Golgi upstream of shear stress. We found that control cells subjected to 48 hours of shear stress exhibited a striking upstream polarization of the Golgi apparatus **(Fig. S5A).** However, siLats1/2 treated cells did not show such a strong polarization **(Fig. S5B)**. This was quantified in sheared conditions by calculating the angle between the vector formed between the EC nuclei and their respective Golgi, and direction of flow **(Fig. S5C).** We found that although the siNT have a seemingly higher proportion of Golgi polarized “against” the flow (∼180° +/- 45° from the flow direction), the siLats1/2 cells exhibited a much wider distribution of angles **(Fig. S5E).** Thus, Lats1/2 signaling is required for correct positioning of the Golgi apparatus under shear stress.

### Lats1/2 mediates VEcad maturation and EC cytoskeletal rearrangements under flow

Another classical endothelial response to blood flow is dynamic changes in cell-cell junctions.^16^ We therefore assessed the status of endothelial VEcad+ junctions in siNT and siLats1/2 cells. We binned VEcad junctions into 3 categories: continuous, serrated, and reticular, based on morphology, as previously described by others^26^ (**Fig. 5H**). We found that in static conditions, all types of junctions were present in control and siLats1/2 cells at about the same frequency (**Fig. 5I, J, M**). Under shear stress, control cells displayed a significant reduction of serrated junctions (mean= 4.84% +/- 5.95% compared to 29.51%+/- 16.91%, p-value= 0.037) and a slight but non-significant increase of continuous junctions (66.675% +/- 13.22% compared to 43.795% +/- 2.877%, p-value= 0.0626) (**Fig. 5K, M**). siLats1/2 ECs, however, did not display any significant differences in any junctional categories after exposure to flow (**Fig. 5L, M**). These results suggest a requirement for Lats1/2 during the dynamic remodeling of vascular cadherin-based junctions under shear stress.

### Endothelial lats1/2 depletion does not cause defects in Akt or KLF2/4 signaling

Activation of signaling pathways such as PI3K^21^ and KLF2/4^19,20,58^ contribute to vessel maintenance and stabilization downstream of laminar shear stress. Therefore, we investigated whether Lats1/2 depletion would cause defects in responses of these signaling pathways to shear stress. We assayed the activation of AKT as a readout for PI3K activation after acute shear stress (30 minutes). Surprisingly, we found that depletion of Lats1/2 with siRNA had no effect on AKT phosphorylation as assessed by Western Blots for pAKT (**Fig. S6A**). Similarly, we assayed *KLF2* and *KLF4* mRNA expression by RT-qPCR and found no significant difference between the activation of either of these two genes **(Fig. S6B)**. Therefore, we conclude Lats1/2 regulation of alignment appears to be independent of the PI3K-AKT and KLF2/4 pathways.

### Loss of Lats1/2 alters flow sensitive pathways

Because previously identified, flow-sensitive signaling pathways did not account for the defects observed in the absence of Lats1/2, we performed bulk RNA-sequencing on the siRNA-treated ECs with and without shear stress for 24 hours to query downstream genetic repercussions. We observed many differentially expressed genes (DEGs) between all conditions observed; of note, there were 707 DEGs in sheared siLats1/2 compared to sheared control cells **(Fig. S6C).** We performed KEGG pathway analysis on all DEGs of siLats1/2 and siNT cells under flow, which revealed a significant difference in pathways such as “Cytoskeleton in muscle cells”, “Hippo signaling pathway-multiple species”, “Cell adhesion molecules”, and “Ras signaling pathway” **(Fig. 6A).** To gain further insight into these differences, we analyzed up- and down-regulated genes separately in siLats1/2 vs. siNT cells under flow **(Fig. 6B)** and in static conditions **(Fig. S6D)**. KEGG pathway “Hippo signaling”, which encapsulates both activators and downstream target genes of both YAP and TAZ was upregulated in sheared conditions, although not significantly different (p-value= 0.053). To gain a better insight into what genes were dysregulated, we generated volcano plots of all DEGs in sheared conditions and queried genes in the respective gene lists of these pathways **(Fig. S6E-H).** Of interest, certain cell adhesion molecules were dysregulated (upregulated: *CLDN11, NECTIN3*, downregulated: *CLDN5, NCAM1*), as were integrins (upregulated: *ITGA4, ITGB8, ITGA11*, downregulated: *ITGA2*), cytoskeletal regulators were upregulated (*MYL9, MYL2*), and placental growth factor (*PGF*), which is known to respond to shear stress^59^ was downregulated in sheared siLats1/2 ECs. These data suggest that there is a shift in the transcriptional landscape of ECs when Lats1/2 are depleted.

**Figure 6.**
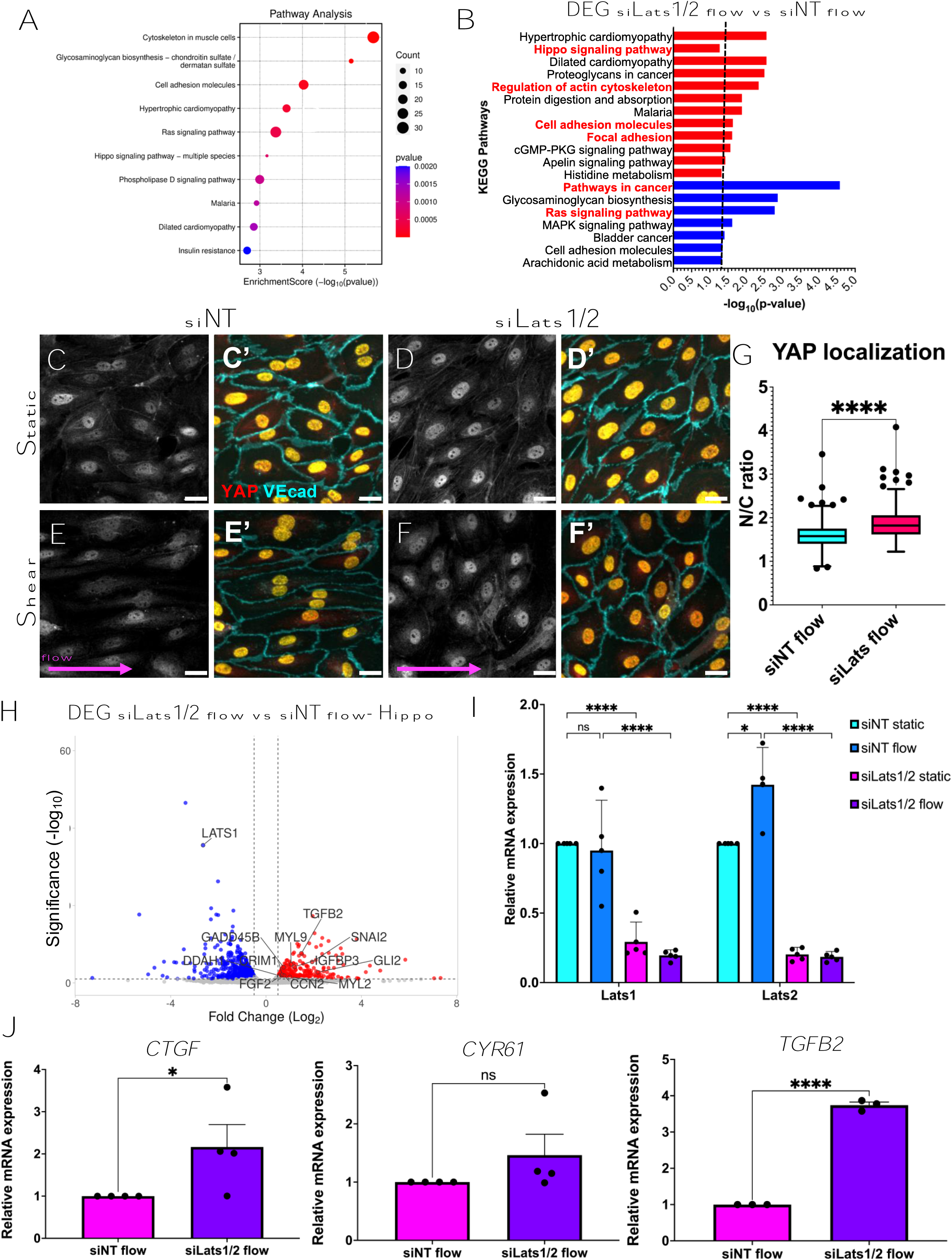
Lats1/2 temporally suppress YAP/TAZ activity under flow. **(A)** Dot plot of KEGG pathway analysis on all DEGs in siLats1/2 flow cells vs. siNT flow cells (24 hours of flow). All p-values are statistically significant (adjusted p-value (FDR) <0.05) and size of dot correlates with number of genes enriched within a pathway. **(B)** Bar chart of selected DEGs in siLats1/2 flow cells vs. siNT flow cells either upregulated (red) or downregulated (blue); x-axis indicates -log(FDR). Dotted line represents significance (FDR<0.05). **(C-F’)** YAP localization in static (**C, C’**) and sheared ECs (48 hours of flow, **D, D’**) with (**D, D’, F, F’**) and without knockdown of Lats1/2 (**C, C’, E, E’**). Pink arrow indicates flow direction. Scale bar= 20 µm. **(G)** Tukey box and whiskers plot of nuclear YAP/ cytoplasmic YAP ratio in sheared conditions. **(H)** Volcano plot of DEGs in siLats1/2 flow cells vs. siNT flow cells. Upregulated (blue) and downregulated (red) genes are color-coded (FDR<0.1, -log2(FC)> 0.5). Labeled genes are knocked down genes (LATS1, LATS2), and selected upregulated Hippo target genes (e.g. *SNAI2, MYL2, CCN2, MYL9, FGF2, IGFBP3, TGFB2).* **(I)** Relative mRNA expression of *Lats1, Lats2* in static and flow in siNT and siLats1/2 conditions using RT-qPCR. All values were normalized to siNT static. Note upregulation of Lats2 under control flow sheared conditions. n= 4 replicates. **(J)** Relative mRNA expression of *CTGF, CYR61*, and *TGFB2* in sheared conditions normalized to siNT flow. There is a significant upregulation of *CTGF* and *TGFB2* in sheared conditions in siLats1/2 cells compared to siNT cells (left, right) but not CYR61. N= 3 replicates. Statistical significance determined by Two-way ANOVA with multiple comparisons (I) or Mann-Whitney U test (G, J). ns= not significant, * indicates p-value<0.05, **** indicates p-value <0.0001.

### Lats1/2 temporally regulate YAP/TAZ activity under flow

Since Lats1/2 cannot cause direct transcription of genes, we asked whether the transcriptional co-activators and Hippo effectors YAP and TAZ were involved. The main function of Lats1/2 are to phosphorylate YAP and TAZ, causing translocation to the cytoplasm and thereby suppressing their transcription activity.^60^ Therefore, we assessed YAP/TAZ localization and activity downstream of flow in the presence or absence of Lats1/2. YAP localization is known to shuttle into the nucleus after acute shear stress and stabilize after sustained or chronic laminar shear stress.^29,30^ Therefore, we hypothesized that Lats1/2 depletion would cause increased nuclear YAP localization. Indeed, siLats1/2 exhibited increased nuclear YAP localization compared to control cells after 24 hours of shear stress exposure **(Fig. 6C-F, G).** Overall, these findings suggest that YAP/TAZ activity is dynamic; nuclear localization of YAP/TAZ spike after acute shear stress, however, Lats1/2 function to reduce YAP/TAZ nuclear localization after sustained shear stress. To better understand how shear stress might impact signaling downstream of Lats1/2 and YAP/TAZ, we assayed the expression of YAP/TAZ target genes in shear stressed siLats1/2 and siNT cells.^61^ We found upregulation of a subset of YAP/TAZ target genes (e.g. *TGFB2, SNAI2, CCN2, IGFBP3, FGF2, MYL9*) **(Fig. 6H)**. To validate our system, we verified knockdown of *LATS1* and *LATS2* under flow via RT-qPCR. Of note, *LATS2* expression was upregulated in control cells under shear stress **(Fig. 6I**, compare blue bars). Additionally, we confirmed sustained expression of target genes *CCN2* (also known as CTGF), *CYR61* and *TGFB2* with qPCR under flow **(Fig. 6J).** These data suggest that Lats1/2 are needed to limit endothelial YAP/TAZ transcriptional activity under shear stress.

### Endothelial Lats1/2 deletion is partially rescued by YAP/TAZ codeletion

Our findings demonstrate that Lats1/2 are necessary for proper EC development *in vivo* and for endothelial cellular responses to shear stress. We next asked whether loss of endothelial Lats1/2 led to hyperactivation of YAP/TAZ signaling, leading to an aberrant response to shear stress and the ultimate demise of the embryos. To test if inactivating YAP/TAZ in the Lats1/2 depleted conditions is sufficient to rescue the defects observed *in vivo* and *in vitro*, we generated *Lats1^ff^;Lats2^ff^;YAP^ff^;TAZ^ff^; Cdh5CreER^T2^* embryos (*Quad^iECKO^*) and evaluated embryos at E10.5. We found that YAP/TAZ inactivation largely abolishes size defects seen in Lats1/2 mice **(Fig. 7A-C)**. Additionally, closer inspection of the yolk sac vasculature revealed that *Quad^iECKO^*embryos had largely normally remodeled vessels, with large and small vessels present in these yolk sacs (**Fig. 7D-F’**). These data suggest Lats1/2 depend on YAP/TAZ activity to cause defects in vascular remodeling and endothelial stability during embryonic development.

**Figure 7.**
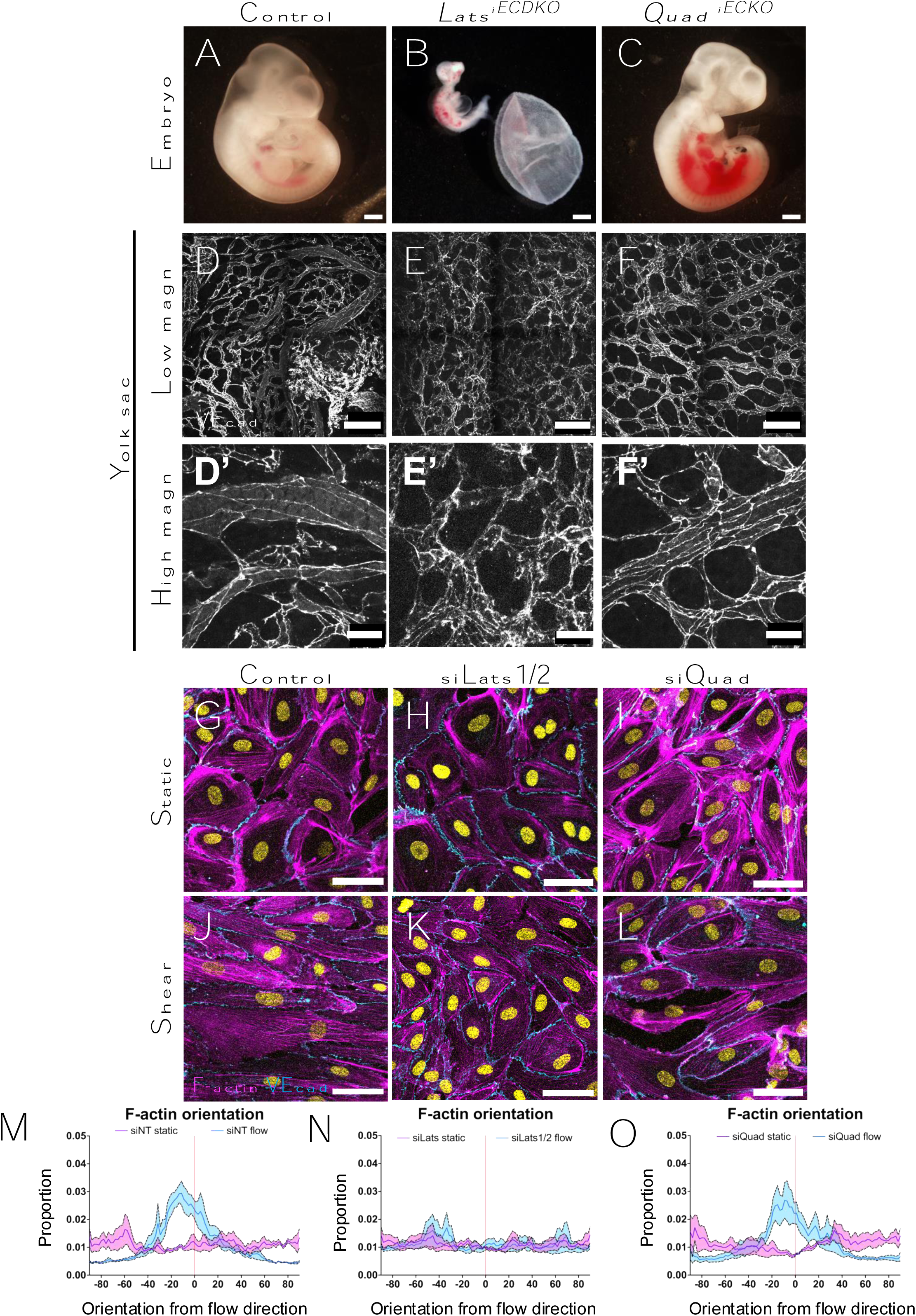
Concurrent deletion of YAP/TAZ in *Lats^iECDKO^* embryos rescues phenotypes *in vivo* and *in vitro.* **(A-C)** Representative brightfield images of E10.5 control (**A**), *Lats^iECDKO^* (**B**), and *Quad^iECKO^* (**C**) embryos. Scale bar= 500 µm. **(D-F)** Representative yolk sacs stained for VEcad in control (**D**), *Lats^iECDKO^* (**E**), and *Quad^iECKO^* (**F**) embryos. Scale bar= 200 µm. (**D’-F’**) Higher magnification of VEcad labeling of respective yolk sacs. Scale bar= 50 µm. **(G-L)** Representative images of F-actin (phalloidin, magenta) orientation in siNT, siLats1/2 and siQuad cells under static (**G, H, I**) and shear stress (**J, K, L**) conditions after 48 hours. **(M-O)** Quantifications of distribution of F-actin stress fiber alignment with respect to flow direction in siNT (**M**), siLats1/2 (**N**), and siQuad (**O**) ECs. Note that siNT and siQuad cells under flow (pink line, M, O), have a strong peak around 0 +/- 30 whereas siLats1/2 cells (pink line, N) do not. Solid line represents average density plot and shaded region represents S.E.M. n = 3 replicates.

To further elucidate if YAP/TAZ transcriptional activity is sufficient to drive vascular remodeling phenotypes, we crossed female mice harboring an active form of TAZ (TAZ-4SA), which is constitutively active due to mutations in the inhibitory Lats phosphorylation sites (TAZ4SA^Tg/Tg^) with an endothelial-specific Cdh5-tTA mouse and evaluated if phenotypes were present at E10.5 (*TAZ4SA^EC^*). *TAZ4SA^EC^*mice were markedly smaller than control embryos, mimicking *Lats^iECDKO^*embryos **(Fig. S7A**, compared to **Fig. 1G**; quantified in **S7E**). Whole mount immunofluorescence of PE revealed a dense, disorganized vasculature in the *TAZ4SA^EC^* mice in the trunk of the embryo, which largely phenocopies *Lats^iECDKO^* **(Fig. S7D,** compared to **Fig. S2J)**. These data indicate that forced nuclear activation of TAZ transcriptional activity is sufficient to drive abnormal EC tube remodeling phenotypes.

Finally, we tested whether Lats1/2 signals function via YAP and TAZ in HPAECs and whether YAP/TAZ knockdown was sufficient to rescue defects seen in response to shear stress in cultured cells. Using siRNA, we concurrently depleted Lats1/2, YAP, and TAZ (siQuad). We then subjected the cells to shear stress for 48 hours and performed immunofluorescence for F-actin using Phalloidin **(Fig. 7G-L)**. We observed that siNT and siQuad cells exposed to LSS elongated along axis of flow **(Fig. 7J, L)**, however, as previously noted, siLats1/2 cells did not elongate to the same extent **(Fig. 7K).** We quantified the proportion of stress fibers that aligned in the direction of flow in these conditions and found that while siLats1/2 cells did not align in the direction of flow **(Fig. 7N),** control and siQuad cells had peaks of aligned fibers close to 0° (within 45°) **(Fig. 7M, O)**. This suggests that Lats1/2 suppress YAP/TAZ activity and subsequent actin reorganization and cell elongation under flow, and that this is required for normal endothelial response to hemodynamic cues.

Overall, these data suggest that Lats1/2 deletion in ECs results in inappropriate upregulation of YAP/TAZ activity. Loss of Lats1/2 blunts the ability of ECs to respond to shear stress from onset of circulation **(Fig. 8)**, leading to a failure of remodeling *in vivo* and *in vitro*. YAP/TAZ deletion in the background of *Lats^iECDKO^* can largely rescue the phenotypes seen both *in vitro* and *in vivo*.

**Figure 8.**
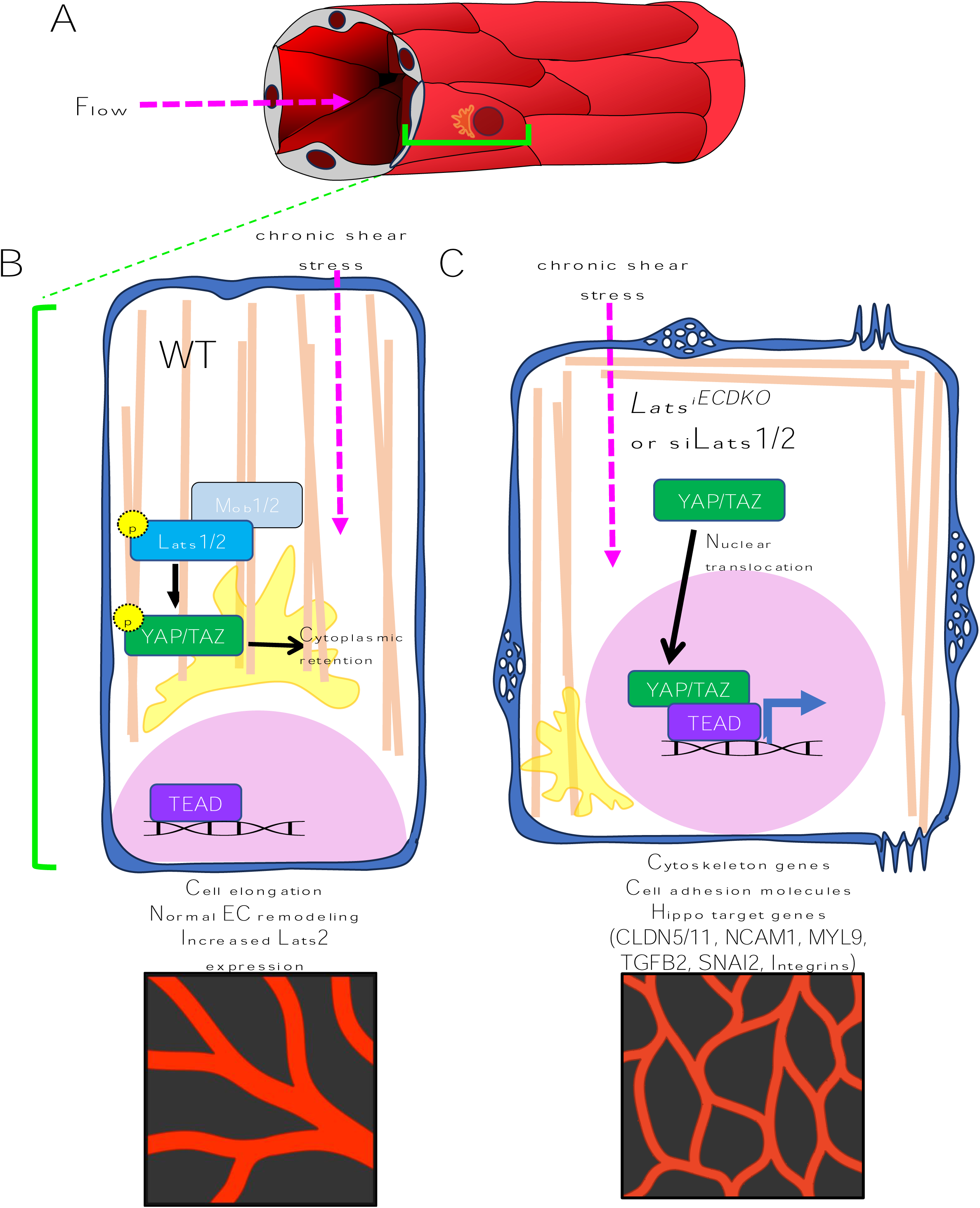
Model – Lats1/2 suppression of Yap is critical for endothelial adaptation to flow. **(A)** Hemodynamic flow along the long axis of blood vessels causes shear stress along the ECs of the vessels, altering their cytoskeleton, junctions, and polarity. **(B)** In the presence of sustained blood flow, ECs in control conditions that have normal Hippo regulation and will facilitate: 1. normal elongation of endothelial cells, 2. Golgi polarization upstream of flow, and 3. remodeling of junctions occurs. (**C**) However, in Lats1/2 depleted conditions, the cytoskeleton is defective, and both Golgi polarization and junctional maturation are defective. Additionally, nuclear YAP/TAZ causes aberrant gene expression of cytoskeletal genes, cell adhesion genes, and Hippo target genes including *CLDN5/11, NCAM1*, *MYL9*, *TGFB2*, and integrins.

## Discussion

In this study, we investigated the role of the Hippo kinases, Lats1 and Lats2 during embryonic blood vessel morphogenesis. We found that mice with endothelial-specific deletion of Lats1/2 display vascular defects at a time when blood vessel remodeling is set in motion by increasing blood flow.^13^ We observe failure of vessel maintenance both in the embryo proper and in the extraembryonic yolk sac, and we find that these occur in a manner dependent on YAP/TAZ signaling. Because of the well characterized role of shear stress on embryonic vascular remodeling in the murine yolk sac and on endothelial lumen maintenance, we tested whether Lats1/2 might mediate the endothelial response to shear stress *in vitro*. Human Pulmonary Artery Endothelial Cells (HPAECs) depleted of Lats1/2 fail to respond to shear stress cues, compared to controls. Lats1/2 depleted cells fail to align and reposition their Golgi. In addition, endothelial cell-cell junction maturation and cytoskeletal reorganization is impaired. We attribute these defects to aberrantly elevated YAP/TAZ signaling, as quadruple deletion of Lats1/2 and YAP/TAZ largely rescues these defects, both *in vivo* and *in vitro*. This study is the first, to our knowledge, to provide an in-depth characterization and phenotypic analysis of Lats1/2 depleted endothelium during embryogenesis. Furthermore, this study places Lats1/2 at the forefront of the molecular machinery transducing the endothelial response to sustained shear stress.

### The Hippo pathway is essential for sustaining vessel assembly and integrity

The proper balance of Hippo signaling has been shown to be critical for tissue development and homeostasis.^62^ In the last few years, proper regulation of the Hippo pathway has also been linked to formation of the vasculature. In 2017, the Koh group identified a requirement for YAP/TAZ in the growth of the postnatal retinal vasculature, as well as in pathological angiogenesis.^27^ They showed that loss of YAP/TAZ resulted in altered cytoskeleton, loss of Cdc42 signaling and loss of filopodia. Furthermore, they showed that loss of Lats1/2 in the postnatal retinal blood vessels led to vessel hyperbranching, and the phenotype could be largely rescued by quadruple knockout of endothelial Lats1, Lats2, YAP, and TAZ. Until this study, there had been no examination of the Lats kinases in forming vasculature. Our work extends analyses of the Hippo pathway to stages when blood vessels first experience hemodynamic flow during development, and we identify Lats1/2 as critical guardians of endothelial stability in part via YAP/TAZ.

Status of YAP/TAZ in ECs also impacts multiple other cellular processes. Loss of YAP and TAZ was found to lead to reduction of proliferation, and conversely, loss of Lats1/2 increased proliferation, and this was shown to occur via the transcription factor MYC ^27^. A second group reported a similar loss of endothelial proliferation when deleting YAP/TAZ in retinal vessels and attributed defects to cytoplasmic YAP/TAZ control of Cdc42 activity.^28^ However, the loss of Lats1/2 also led to a decrease in Cdc42 activity, as phosphorylated YAP/TAZ in the cytoplasm is required. In our system, however, we found that EC proliferation was not elevated at the early timepoints we examined. Rather, proliferation was slightly reduced. Similarly, we did not find a difference in proliferative capacity of cultured ECs with depleted Lats1/2 activity at various time points following siRNA transfection. We note this with interest, as extrinsic forces such as shear stress can cause dynamic changes in EC proliferation.^53^

Instead, we found that loss of Lats1/2 resulted in marked changes in endothelial cell-cell junctions. We observe striking junctional defects in vessels of the embryo, and in those in the remodeling yolk sac. Our findings are reminiscent of work from the Gerhardt group who showed that loss of YAP/TAZ led to downregulation of EC adherens junction turnover, loss of cell-cell adhesion maturation and cell shape alteration.^26^ These defects could be rescued in part by suppression of BMP and Notch signaling. The role of Lats1/2 in this model system was not addressed in this study, and it therefore remains unclear if YAP/TAZ activity was dependent or independent of Hippo signaling. A critical additional finding is that in our HPAEC system, we did not detect significant differences in VEcad junction morphology until cells were exposed to shear stress. Hence it is possible that junctional maturation requires presence of YAP/TAZ regardless of biomechanical forces from blood flow, however, elevated levels of YAP/TAZ due to Lats1/2 depletion are deleterious to junctional maturation when challenged by hemodynamic stress. These studies together point the importance of the Hippo pathway to endothelial cell biology and vessel integrity.

### Vascular defects in the absence of Lats1/2 begin at the onset of hemodynamic flow

It is striking that the timing of the embryonic defects we observe in the absence of Lats1/2 is coincident with the timing of remodeling of the initial embryonic vasculature, a process that is highly dependent on blood flow.^12^ Vessels in our mutants begin to fail around E9.0, at a time when the heart beats progressively more strongly, and blood circulation begins. We postulate that the loss of Lats1/2 leads to the demise of the embryo due to failure of junction remodeling and cell shape changes when circulation ramps up and puts pressure on the endothelium. These failures are due to the inability of ECs to sense flow, and structurally they lead to lumen collapse, formation of occlusions, and compromised circulation.

We used *in vitro* modeling to test the hypothesis that Lats1/2 transduces flow adaptation, and we show that ECs lose the ability to respond to shear stress cues upon Lats1/2 depletion. However, there are important caveats revealed by our different experiments. Case in point, for example, is that blocking Lats1/2 activity in 3D EC tube networks in static conditions results in failure of vessel maintenance. Treatment with the Lats1/2 inhibitor TRULI after the formation and establishment of tubes *in vitro* results in EC tube regression and collapse. This points to a requirement for Lats1/2 activity that is flow independent. Or, it points to TRULI targeting pathways we are as yet unaware of, beyond Lats1/2.^63^ By contrast, ECs treated with siRNA targeting Lats1/2 when plated on plastic in static conditions, did not display major changes in junctional organization, cell shape, or proliferation, even though we detected significant shifts in the transcriptional landscape in static conditions.

This brings up the question: what is the requirement for Lats1/2 in ECs in either static or under hemodynamic flow forces, either in soft (embryo or collagen matrices) versus hard (plastic from petri dish) surfaces? We speculate that ECs in 3D matrices are more likely to reveal failures of tube formation that would not be detectable in 2D cultures on plastic. Our previous work underscored how cytoskeletal signaling failures displayed different cellular defects in different environmental conditions.^47^ We speculate that the loss of Lats1/2 results in loss of sensing of blood flow, but that other cell-intrinsic, functional changes unrelated to the sensing of shear stress are also dependent on Hippo signaling. Nevertheless, the changes we see lead to an ultimate failure of the functional response of endothelial cells: their ability to respond to shear stress.

### Are Lats1/2 required for endothelial lumenogenesis?

It is also important to note that although vasculogenesis occurs normally in our model system, the Lats1/2 deletion carried out by the VE-cadCreERT2 (Cdh5(*PAC*)-*CreERT2*) allele,^35^ Lats1/2 may play a role embryonic lumenogenesis. Our lab previously characterized the timing of deletion using other endothelial drivers, such as Tie2Cre, and we showed available endothelial Cre drivers do not delete the floxed gene of interest until after lumenogenesis occurs.^46^ We attempted to address this by looking at tomato expression in E8.5 embryos and did see tomato expression at this stage. However, the removal of proteins and the induction of recombination do not always coincide ^64^. This limitation of our system must be considered when evaluating the role of Lats1/2 in the early vasculature. Lats1/2 may play a yet unappreciated role in formation of the dorsal aortae and drivers such as CAGGCreER^TM 65^ or another early and ubiquitously expressed inducible driver could help decipher earlier roles for Lats1/2. Nevertheless, the requirement for Lats1/2 during vascular remodeling, which occurs after the onset of flow, is the primary defect we observe using our present model system.

### Suppression of YAP/TAZ by Lats1/2 is required by ECs to sense flow

Our findings identify a novel requirement for restraint of YAP/TAZ signaling by Lats1/2 in the adaptation of laminar shear stress by ECs. Others have identified a role for YAP/TAZ localization and activity during the onset of shear stress in zebrafish and cultured ECs.^29^ However, YAP/TAZ are generally cytoplasmic in mature, quiescent vessels, which are under sustained laminar shear stress.^30^ We identify a novel role for the upstream regulators Lats1 and Lats2 in modulation of YAP/TAZ in developing vessels. Our study did not test the phosphorylation of Lats1/2 or their interaction with YAP/TAZ under sustained flow. However, we show that Lats2 expression in our system is upregulated after 24 hours of shear stress. Lats2 may be upregulated over time as a negative regulator of YAP/TAZ under shear stress. This is supported by evidence in MCF10A cells, where overexpressing TAZ4SA in these cells leads to upregulation of endogenous protein levels of Lats2 as well as phosphorylation of Lats1/2, leading to negative regulation of YAP/TAZ activity.^66^ Therefore, we propose that Lats2 activity under shear stress acts to dampen YAP/TAZ activity under sustained shear stress, at a time where YAP/TAZ activity is no longer needed, and cells instead need to enter a quiescent state as vessels differentiate. Lats1/2 could act as modulators of YAP/TAZ target gene expression in a negative feedback loop, and deletion of Lats1/2 leads to deleterious effects due to aberrantly sustained YAP/TAZ activity.

### Narrow range of Hippo signaling tolerated in ECs

A previous study reported that EC-specific deletion of YAP/TAZ also leads to embryonic angiogenesis defects and lethality in a VEGF-dependent way.^25^ How is it, then, that both YAP/TAZ deletion and YAP/TAZ hyperactivation by Lats1/2 deletion can both cause lethality? Interestingly, previous studies have argued that YAP/TAZ activation could be due to VEGF/ VEGFR2 feedback inhibition of Hippo signaling (e.g. inhibition of Lats1/2) *in vitro* and *in vivo* to regulate the endothelial barrier.^27,67^ However, YAP/TAZ activity is dispensable for barrier integrity during adulthood.^27^ The complexity of signaling inputs on endothelium, however, is underscored by the Koh group’s finding that VEGFC activates the Hippo pathway, which is opposite of what VEGFA does in blood ECs *in vitro*. Additionally, VEGF-VEGFR activity and shear stress are known to have competing roles in the developing retina. Therefore, it is possible that YAP/TAZ and Lats1/2 are needed at different times and in different contexts. However, loss of either one throws off the balance of endothelial integrity, leading to loss of vascular resilience and embryonic lethality.

### Independent roles of Lats1/2 and YAP/TAZ

Understanding whether Lats1/2 and YAP/TAZ function in a linear signaling relationship in ECs will be key to future development of any pro- or anti-angiogenic clinical approaches. In our model, although the embryo defects in *Quad^iECDKO^* embryos were largely rescued compared to *Lats^iECDKO^* embryos, about half of the *Quad^iECDKO^* embryos still experienced hemorrhaging. Of note, in different contexts, Lats1/2 have roles independent of YAP/TAZ signaling, such as mitosis, apoptosis, and actin polymerization.^68^ There is a possibility that the incomplete rescue of defects in both our *in vivo* and *in vitro* Lats deletion models could be due to non-canonical Lats1/2 roles outside of their characterized role in YAP/TAZ localization and transcription. Intriguingly, Lats1 and Lats2 have been placed at the membrane in other cell types.^69^ It is possible they interact with junctional molecules after the onset of shear stress upstream and act in ways completely independent of YAP/TAZ signaling. Additionally, the regulation of translocation and downstream signaling of YAP/TAZ in many cells *in vivo* or *in vitro* have been debated. Some studies have shown YAP/TAZ translocation can be independent of canonical regulation by Lats1/2, but rather dependent on F-actin dynamics instead.^23^ As mentioned before, YAP/TAZ depletion in embryonic ECs also causes defects in and of itself, since some level of transcriptional activity from YAP/TAZ are required during vascular development. The incomplete rescue we observe in our quadruple knockout/knockdowns could be due to the roles YAP/TAZ play on their own, and co-depletion of YAP/TAZ with Lats1/2 could cause phenotypes similar to YAP/TAZ depletion alone. Teasing out the temporal and spatial regulation of Hippo signaling and understanding how it cooperates with other pathways in endothelium, will propel our knowledge of this pathway in vascular morphogenesis and will facilitate the identification of new targets to treat vascular diseases.

### Summary

This study provides insight into a previously underappreciated role for Lats1/2 in endothelial mechanotransduction and development of embryonic blood vessels. Disruption of fluid forces or downstream pathways can lead to a multitude of diseases and conditions, such as atherosclerosis, cancer, and arteriovenous malformations. Thus, studying how molecules regulate the process of EC maintenance and stability downstream of mechanical forces will be critical to the development of treatments for vascular diseases.

## Acknowledgements

We thank the entire Cleaver lab for critical discussions and input on the manuscript. We thank Randy Johnson for the *Lats1^flox^; Lats2^flox^* mice, Eric Olson for the *Yap^flox^;Taz^flox^*mice, and Duojia Pan for the *TRE-TAZ4SA* mice. We thank the UTSW Genomics Core for processing, sequencing, and initial data analysis of RNA for bulk RNA-seq. We thank present and past Cleaver lab members for valuable input and discussion from the inception of this work and assistance with tissue culture (Anne Ryan, Caitlin Maynard, Neha Ahuja, Stephen Spurgin, Luis Rangel, Peter Luo).

## Footnotes

## Author contributions

Conceptualization: M.A.C., O.C.; Methodology: M.A.C., O.C.; Software: M.A.C; Validation: M.A.C.; Formal analysis: M.A.C., O.C.; Investigation: M.A.C., T.P., Y.F, S.R.M.A., A.M., O.C.; Resources: O.C.; Data curation: M.A.C.; Writing - original draft: M.A.C., O.C.; Writing - review & editing: M.A.C., T.P., A.M., V.D.V., G.E.D., O.C.; Visualization: M.A.C.; Supervision: O.C.; Project administration: O.C.; Funding acquisition: O.C.

## Funding

This research was supported in part by grants from the National Institute of Diabetes and Digestive and Kidney Diseases (DK106743, DK079862 to O.C.;); the National Heart Lung and Blood Institute (HL113498 to O.C.); the National Academy of Science, Engineering, and Medicine Ford Foundation Dissertation Fellowship (M.A.C.); and the Foundation Leducq grant (21CVD03 to O.C.). Open Access funding provided by University of Texas Southwestern Medical Center. Deposited in PMC for immediate release.

## Data availability

RNA-seq data have been deposited in the Gene Expression Omnibus (GEO) under the accession number xxx.

---

**Figure S1.**
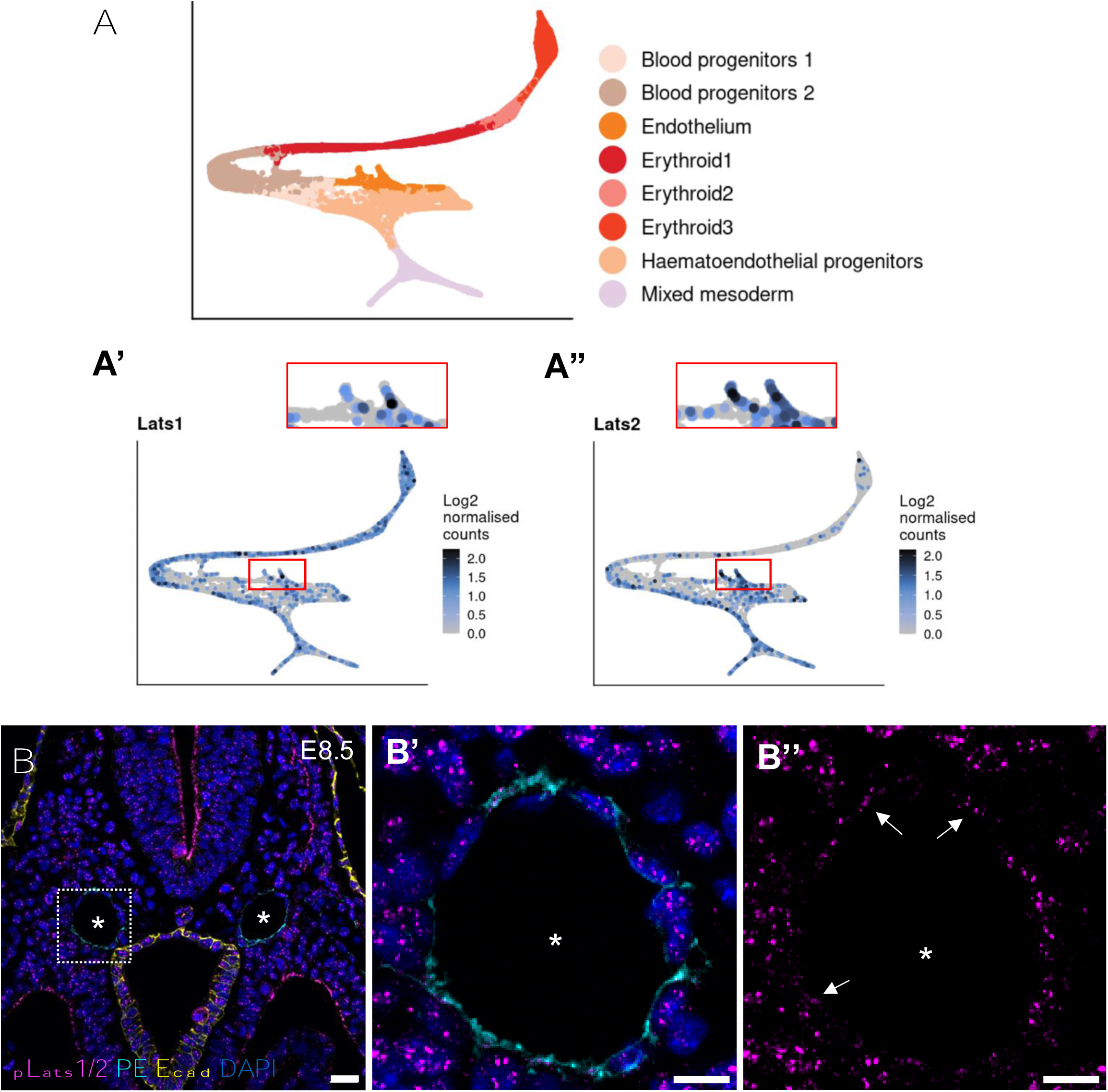
Presence of LATS1/2 in embryonic endothelium.

**Figure S2.**
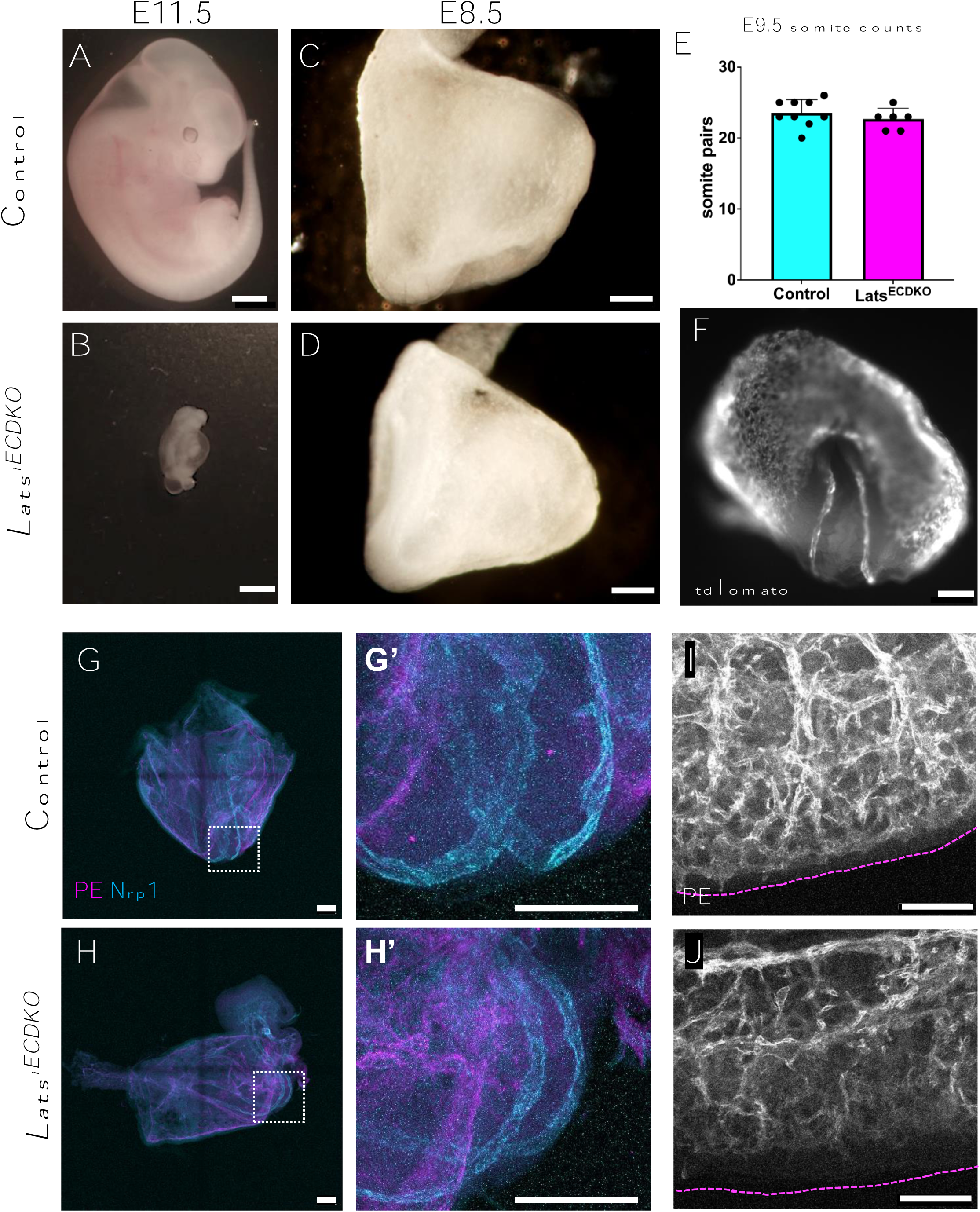
Loss of Lats1/2 activity in endothelium leads to embryonic lethality and to tubule collapse *in vitro*. **(A-B)** Representative brightfield images of control (A) and *Lats^iECDKO^* (B) embryos at E11.5. Note opaque appearance of *Lats^iECDKO^* embryo. n> 3 embryos/ genotype. Scale bar= 1mm **(C-D)** Representative brightfield images of E8.5 control (C) and *Lats^iECDKO^* (D) embryos. Scale bar=250 µm. **(F)** Somite counts of E9.5 control and *Lats^iECDKO^* embryos. n= 6-9 embryos/ genotype. **(E)** Endogenous tdTomato expression of *Lats^iECDKO^* embryo at E8.5 during dissection. Scale bar= 150 µm. **(G,H)** Expression of arterial marker Neuropilin1 (Nrp1) in E8.5 control and *Lats^iECDKO^* embryos. **(G’, H’)** magnification of G and H. Scale bars= 150 µm. **(I, J)** Whole mount immunolabeling of PECAM/ Emcn of trunk region of E9.5 control and *Lats^iECDKO^* embryos. Dotted line represents extent of dorsal midline. Scale bar= 100 µm.

**Figure S3.**
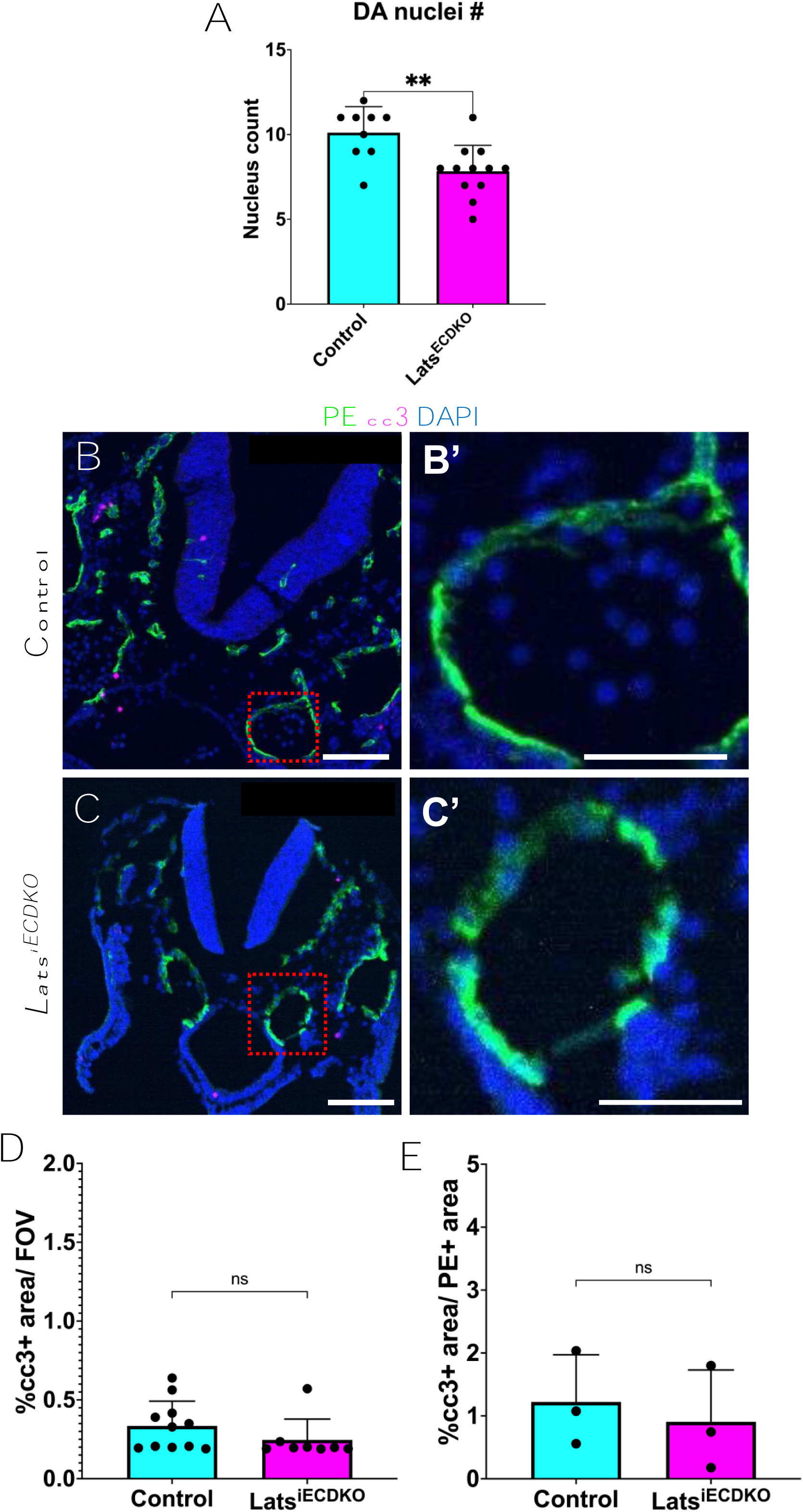
Decreased dorsal aortae ECs in *Lats^iECDKO^* embryos are not due to apoptosis. **(A)** Quantification of nuclei present in aortic ECs in E9.5 control and *Lats^iECDKO^* embryos. Each dot represents one cross section through the aorta; n= 3 embryos/ genotype. **(B, C)** Relatively low abundance of cleaved caspase 3 (cc3, magenta) in immunostained control and *Lats^iECDKO^* embryos. Scale bar= 200 µm. **(B’ C’)** Enlarged boxed regions from B and C respectively. Scale bar= 50 µm. **(D)** Quantification of cc3+ area/ FOV. Each value represents measurement from one image. Representative of 3 embryos mutants and two stage matched controls. **(E)** Quantification of cc3+ area in endothelial cells only (PE+ area). Each value represents average from multiple FOVs/ embryo. Statistical significance determined by unpaired t-test (A) and Mann-Whitney U test (D, E). ns= not significant; ** indicates significance at p< 0.01.

**Figure S4.**
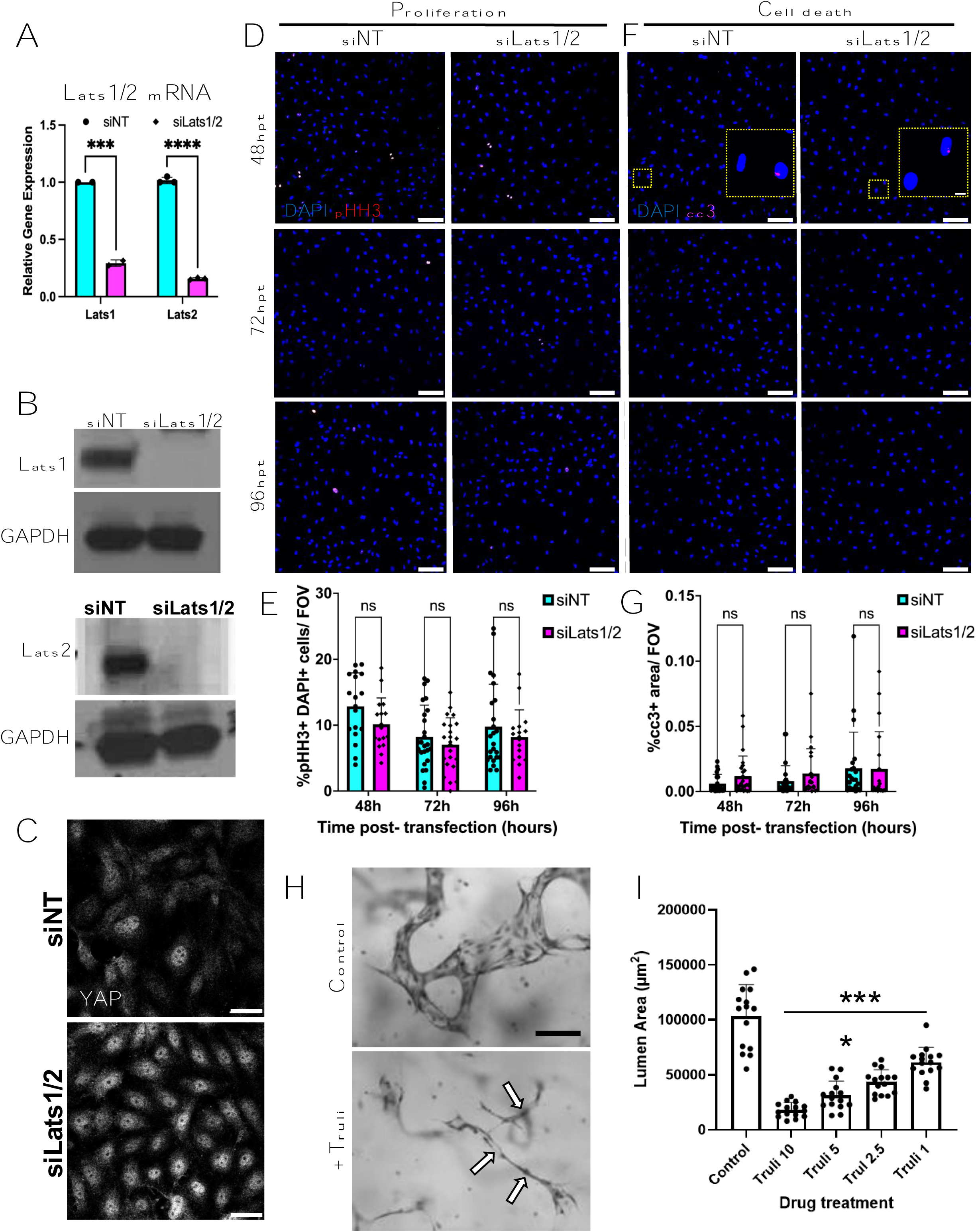
Knockdown of Lats1/2 via siRNA treatment results in Yap activation, but no changes in cell proliferation. **(A)** Verification of knockdown of mRNA of Lats1 and Lats2 48 hours post siRNA transfection in HPAECs using RT-qPCR. n= 3 **(B)** Verification of knockdown of protein of Lats1 (left) and Lats2 (right) 72 hours post siRNA transfection in HPAECs. **(C)** YAP immunostaining in monolayers of HPAECs 72 hours post siRNA transfection. siLats1/2 cells have visibly higher nuclear YAP levels. Scale bar= 50 µm. Representative of at least 3 separate experiments. **(D)** Immunofluorescence images of pHH3 (orange) in siNT and siLats1/2 cells 48-, 72-, and 96-hours post siRNA transfection (hpt). Representative of 3 separate experiments. Scale bar= 100 µm. Quantified in **(E).** Bar represents mean +/- SD. Each value represents one image analyzed per replicate. n= 16-24 20X FOVs; 3 replicates. **(F)** Immunofluorescence images of cleaved-caspase 3 (cc3) (magenta) in siNT and siLats1/2 cells 48-, 72-, and 96-hours post siRNA transfection (hpt). Inset image is magnified example of cc3+ cell. Representative of 3 separate experiments. Scale bar= 100 µm. Quantified in **(G).** Bar represents mean +/- SD. Each value represents one image analyzed per replicate. n= 19-24 20X FOVs; 3 replicates. **(H)** ECs were seeded in 3D collagen gels for 48 hr to allow for the formation of tube networks, and then the Lats1/2 inhibitor, TRULI was added at varying concentrations vs. control. At 48 hr, EC tube networks were treated under control conditions or TRULI was added at 5 µM. After 48 hr, cultures were fixed, stained, and photographed. Arrows indicate collapsed EC tubes. Scale bar= 100 µm. **(I)** At 48 hr, EC tube networks were treated with the following TRULI concentrations (10, 5, 2.5, 1 µM) vs. control. After 48 hr, cultures were fixed, stained, photographed, and quantitated for EC lumen area. **** indicates significance at p< 0.0001 compared to control. Statistical significance determined by Mixed effects analysis with multiple comparisons (E, G). ns= not significant.

**Figure S5.**
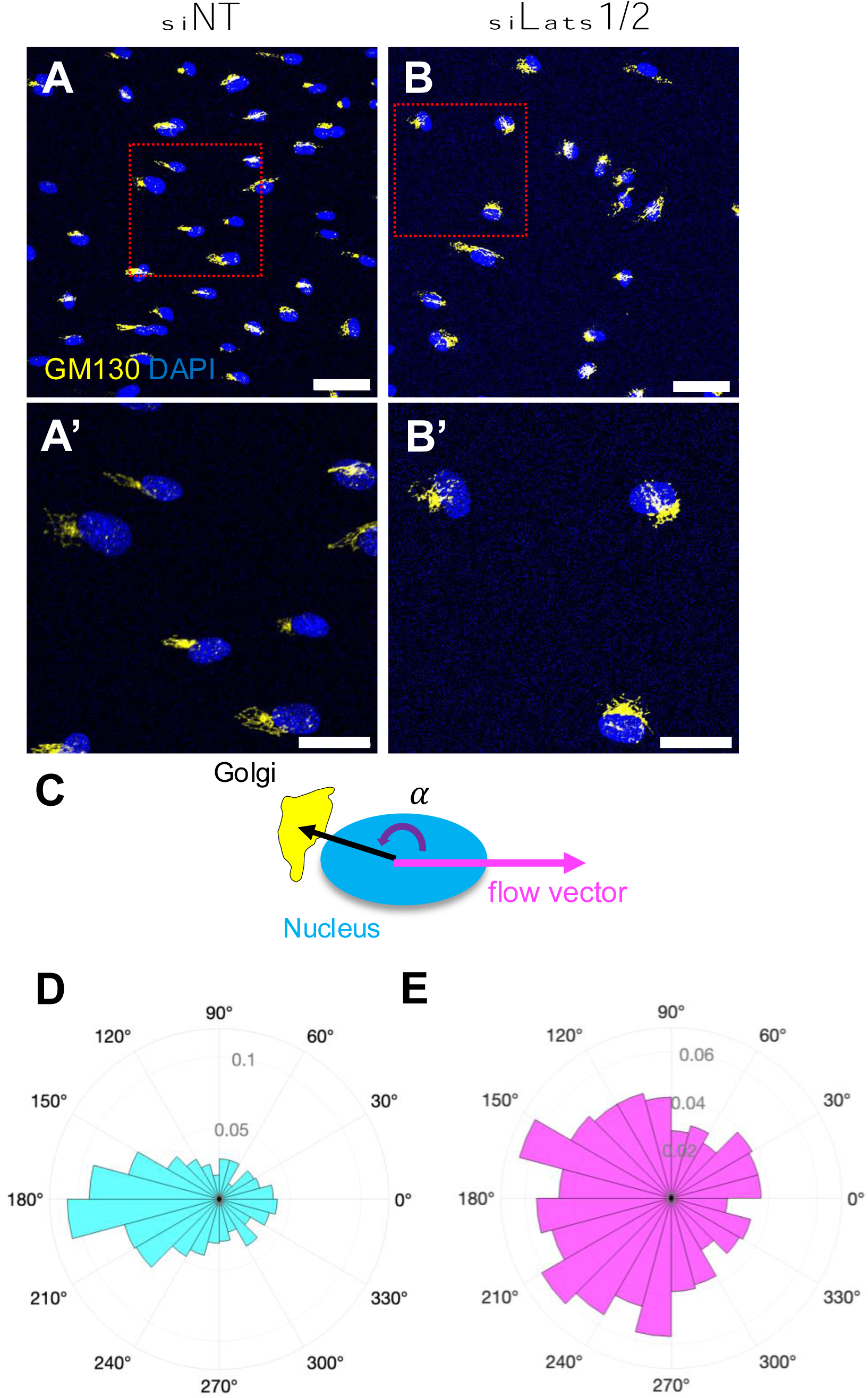
GM130 polarity under flow is attenuated in Lats depleted cells. **(A-B)** Representative images of Golgi (GM130, yellow) polarization with respect to nucleus (DAPI, blue). Scale bar= 100 µm. (A’-B’) magnified boxed regions indicated in A and B respectively. Scale bar= 50 µm. **(C)** Diagram representing calculations used for quantifications shown in D-E. Black arrow indicates vector from centroid of nucleus to centroid of Golgi. Pink arrow indicates flow direction. Purple curved arrow represents calculated angle (α). **(D, E)** polar histograms of nucleus-Golgi vectors with respect to flow direction as depicted in C. siNT cells exhibit a strong polarization “against” flow (180 +/- 30) whereas siLats1/2 are more evenly distributed across all angles. Graph represents n= 3 separate experiments.

**Figure S6.**
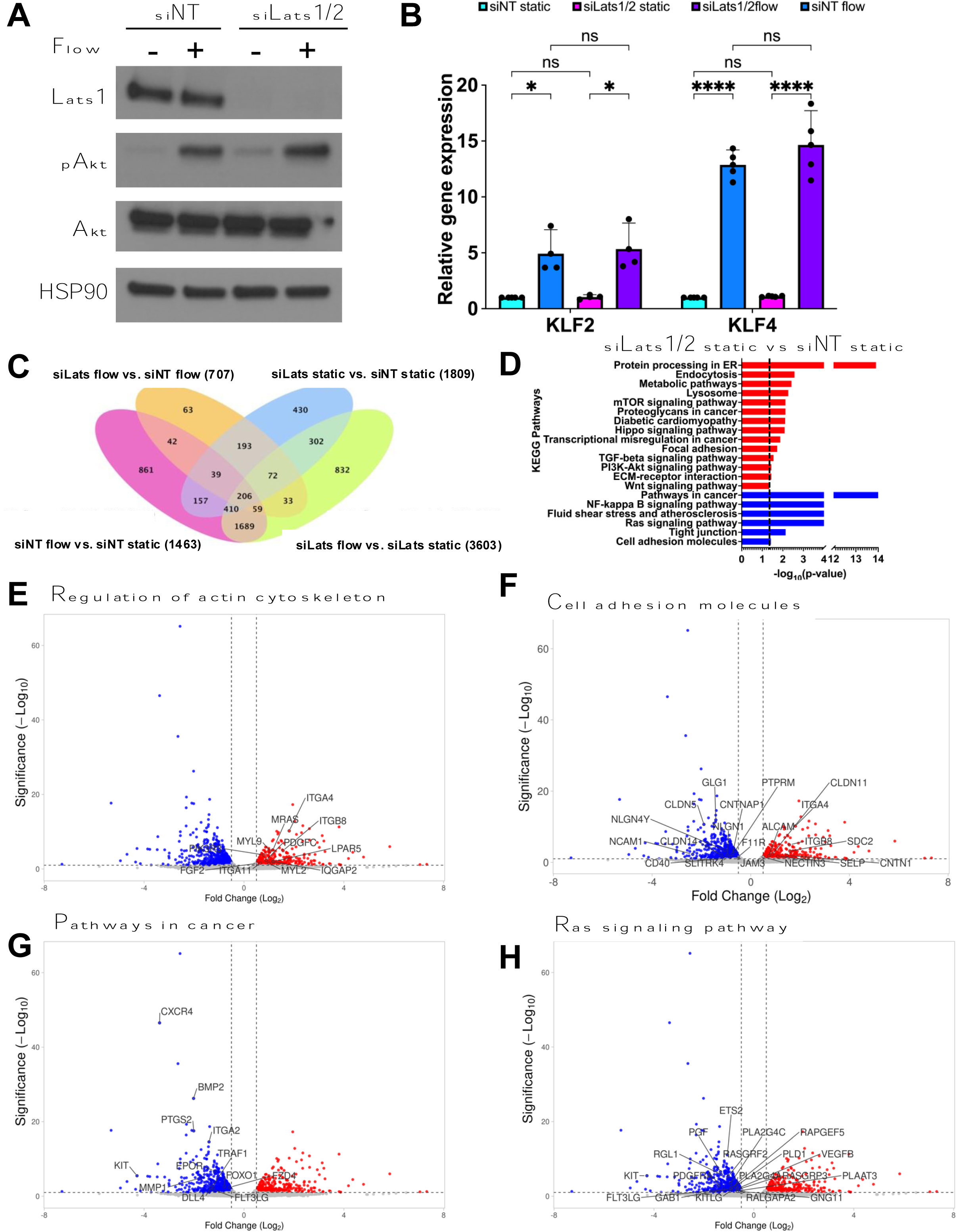
Lats1/2 depleted cells have altered transcriptomic response to flow, but do not affect pAKT or KLF2/4 responses. **(A)** Western blot on confluent monolayer showing upregulation of pAKT under shear stress in siNT and siLats1/2 cells after exposure to 30 minutes of laminar shear stress. Representative of 2 separate experiments. **(B)** *KLF2* and *KLF4* gene expression analysis in siNT and siLats1/2 cells after exposure to shear stress. n= 4 separate experiments for *KLF4* and 3 for *KLF2*. **(C)** Venn diagram of all DEGs in siNT and siLats1/2 cells in static vs. flow conditions. Numbers in parentheses indicate number of DEGs in each condition. Genes were considered significant if they had an FDR <0.1 **(D)** Bar chart of selected KEGG pathways made from genes upregulated (red) and downregulated (blue) in siLats1/2 static vs. siNT static cells. Dotted line indicates significance (FDR< 0.05). **(E-H)** Volcano plots of DEGs between siLats1/2 vs. siNT flow conditions. Labeled genes are terms in bolded KEGG terms from Fig. 7G. **(E)** Upregulated: “Regulation of actin cytoskeleton” (e.g. ITGA4, ITGB8, LPAR5, IQGAP2, MYL2, MYL9, MRAS, FGF2, ITGA11, PDGFC); **(F)** upregulated and downregulated: “Cell adhesion molecules” (e.g. down: CLDN5 CLDN14, CD40, up: ITGA4, CLDN11, CNTN1. **(G)** downregulated: “Pathways in cancer” (e.g. CXCR4, BMP2, ITGA2, PTGS2) and **(H)** “Ras signaling pathway” (e.g. KIT, PGF, ETS2, RAPGEF5). Statistical significance determined by 2-way ANOVA with multiple comparisons (B). ns= not significant,* indicates p-value< 0.05, **** indicates p-value< 0.01.

**Figure S7.**
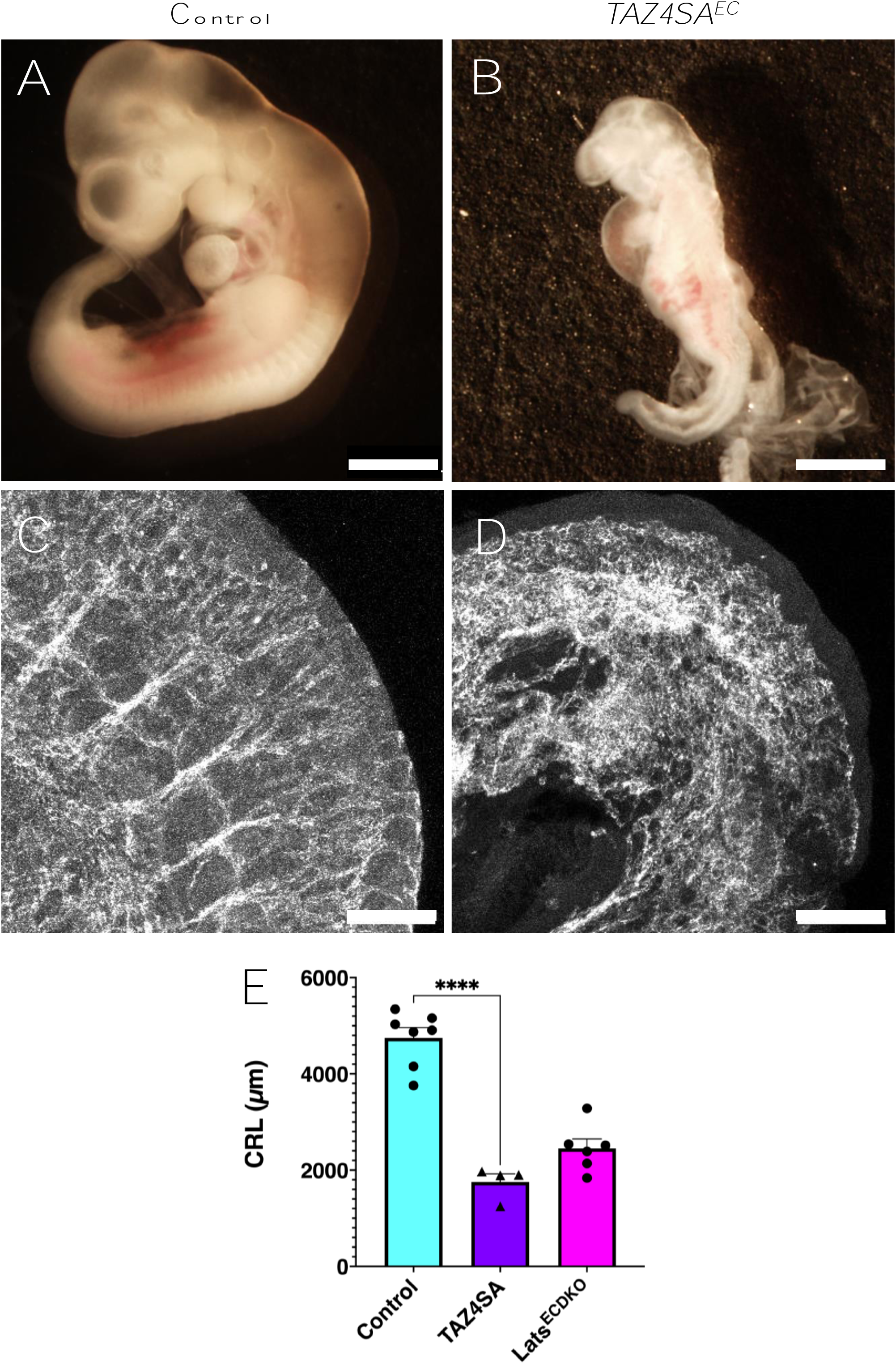
Hyperactivated nuclear TAZ is sufficient to induce vascular remodeling defects. **(A-B)** Representative brightfield images of E10.5 control **(A)** and TAZ4SA^EC^ **(B)** embryos. Scale bar= 500 µm. **(C-D)** Representative images of VEcad staining in trunk vessels of control (C) and TAZ4SA^EC^ (D) embryos at E10.5. Scale bar= 100 µm. n= 2 embryos/ genotype. **(E)** Quantification of embryo crown to rump length (CRL). Each value represents one embryo. Statistical significance determined by one-way ANOVA with multiple comparisons. n=4-7 embryos/ genotype. **** indicates p-value <0.0001.

## Notes

### Competing Interest Statement

The authors have declared no competing interest.

